# Epigenetic Clocks Uncover the Natural History of Ageing in Wild Mice

**DOI:** 10.1101/2025.04.09.648041

**Authors:** Sarah E. Wolf, Simon A. Babayan, Riccardo E. Marioni, Steve Horvath, Amy B. Pedersen, Tom J. Little

## Abstract

Environmental stressors are expected to accelerate senescence but demonstrating this requires both a measure of senescence and knowledge of chronological age - data rarely available for most wild animals. Here, we constructed a DNA methylation-based epigenetic clock for the wood mouse (*Apodemus sylvaticus*) using known-aged laboratory individuals and applied it to a wild population of unknown ages. This represents one of the first demonstrations that a laboratory-trained epigenetic clock can be used to infer age structure in free-living animals. Multiple lines of evidence validated use of the clock in wild individuals: age estimates fell within expected ranges, recapitulated population demographic structure, and recovered predicted signatures of senescence, including an increase in infection probability with age.

Moreover, wild mice given experimental food supplementation and parasite-removal drugs exhibited slower epigenetic ageing rates. Together, our findings demonstrate that epigenetic clocks can infer both chronological age and ageing rates in the wild, providing a powerful framework to identify the ecological and individual determinants of ageing.

## 1. INTRODUCTION

Accurate age estimation in wild animals can unlock powerful insights into ecological and evolutionary patterns. On the one hand, age is a key demographic parameter linking individual life histories to population-level outcomes; on the other hand, variation in ageing rates over a lifetime can reveal fundamental mechanisms underpinning the evolution of senescence. However, age structure data are often difficult to obtain for species that are inaccessible during or shortly after birth. Most indirect estimates of age lack both generality and accuracy. For example, body mass-based estimates are confounded by variation in condition, while others – such as otolith growth rings, eye lens weight, or bone histology (Esteban et al., 1996; Pannella, 1971; Rowe et al., 1985) – lack generality and require post-mortem sampling. Estimates of exact age via telomere length are poor in wild animals (Pepke, 2024).

By contrast, epigenetic clocks appear to provide very accurate age estimates based on patterns of DNA methylation (DNAm). Essentially, DNAm at a subset of cytosine-guanine sequences (CpGs) changes in a predictable ‘clock-like’ manner with age (Teschendorff & Horvath, 2025). DNAm can modulate gene expression (Razin & Kantor, 2005), thus providing a mechanistic link between molecular ageing and phenotypic outcomes. Epigenetic clocks have been trained to predict chronological age and age acceleration (Le Clercq et al., 2023; Lu et al., 2023), both of which have been linked to various degrees to elevated disease risk and mortality in humans (Marioni et al., 2016; Perna et al., 2016). While some clocks are species- or tissue-specific (K. J. Peters et al., 2023; Zoller et al., 2024), others show remarkable generality. For instance, the Pan-Mammalian clock (Lu et al., 2023) accurately estimates age across 185 mammals, underscoring the evolutionary conservation of age-related DNAm patterns. More broadly, that DNA methylation is present across vertebrates makes this a powerful approach for uncovering general principles of ageing.

Pioneering work on epigenetic clocks was conducted primarily in humans and laboratory systems (Horvath & Raj, 2018). While these studies provide foundational knowledge on the mechanisms regulating senescence (López-Otín et al., 2023), they do not provide insight into the genetic, physiological, and environmental variation characteristic of natural populations. Over the past decade, evidence has shown that DNA methylation levels respond to natural stressors (e.g., Bentz et al., 2021; Sepers et al., 2021; Watson et al., 2021), but only recently have epigenetic clocks been applied to wild animals, primarily to build clocks that predict chronological age and to identify the biological functions associated with age-related CpG sites (Le Clercq et al., 2023; Lu et al., 2023). A smaller number of studies have begun to explore how the environment contributes to overestimation of chronological age, or “age acceleration”, such as high-ranking social status (Anderson et al., 2021), reduced sibling number (Haller et al., 2025), life outside of hibernation (Pinho et al., 2022), sedentary lifestyle (Cristofari et al., 2026), and reintroduction of laboratory mice to wild habitats (Zipple et al., 2025).

Assessing epigenetic age acceleration in wild populations extends insights from laboratory models to ecologically relevant contexts. The wood mouse (*Apodemus sylvaticus*) offers a valuable wild analogue to laboratory mice: short-lived (≤ 1 year) and amenable to repeatedly sampling, yet true age is difficult to determine because individuals are not captured until weeks after birth. They are particularly valuable because wild wood mice experience multiple, concurrent stressors, including food limitation and parasite infection. In humans, both are associated with accelerated epigenetic ageing (Durso et al., 2022; Galkin et al., 2023; Gupta & Vadde, 2026), and many clocks include CpGs linked to growth, metabolism, and immune function (Horvath & Raj, 2018). The impact of a stressor on epigenetic age depends on the functional composition of the clock: a clock biased towards metabolism-associated CpGs may be most sensitive to energetic stress. Because both food limitation and infection constrain resources, epigenetic age may respond to the stronger stressor or both additively. These scenarios suggest that environmental factors can shape DNAm and influence biological ageing. Applying epigenetic clocks to wood mice presents an opportunity to expand our understanding of ageing beyond larger, long-lived species where most epigenetic ageing work has been carried out.

Here, we develop a highly accurate epigenetic clock for the wood mouse using laboratory-reared individuals, providing a practical solution for ageing wild systems lacking known-age data. We complement this with an epigenome-wide association study and GREAT analysis to identify the biological functions associated with chronological age. We then apply this epigenetic clock to wild individuals of unknown age, demonstrating that it can resolve the age structure of a previously “un-ageable” population. We validate the clock using multiple independent predictions of age-related change, confirming its reliability and ecological relevance in wild populations. Finally, by tracking within-individual changes in epigenetic age, we test whether reducing environmental stress - via food supplementation and parasite removal, slows epigenetic ageing in the wild. Altogether, our findings showcase a major step forward for epigenetic clocks, enabling precise estimates of age and ageing in the wild and illuminating the ecological and individual factors shaping senescence.

## 2. MATERIALS and METHODS

In order to assess our three major aims, we collected DNA methylation (DNAm) data from (1) known-age captive wood mice and (2) unknown-age wild wood mice. All animal work was performed under UK Home Office PPL PP4913586 and with permission from landowners, in compliance with the Animals (Scientific Procedures) Act of 1986 and approved by the University of Edinburgh Ethical Review Committee.

### 2.1 Sample Collection from known-age captive wood mice

DNA samples for methylation analyses and clock construction were collected from known-age wood mice that were bred and maintained at the University of Edinburgh, UK, in our wild-derived, but laboratory-reared, outbred colony. Mice were housed under standard 12 h light: 12 h dark conditions with *ad libitum* access to food and water. Mice were not subject to any interventions before sampling. Ear biopsies were collected and stored at -80°C until further processing.

In total, we collected 129 ear samples from mice aged 178.8 ± 11.4 days (25 – 383 days, n = 65 males, n = 64 females). DNAm data was generated in three batches: (1) 12 mice aged 194 ± 29.3 days (77 – 383 days, n = 5 males, n = 7 females); (2) 86 mice aged 176 ± 15.8 days (25 – 324 days, n = 46 males, n = 40 females); and (3) 31 mice aged 179 ± 14.8 days (59 – 382 days, n = 14 males, n = 17 females). While the 1^st^ and 3^rd^ batches contain single samples per mouse, the 2^nd^ batch contains two ear biopsies per mouse: at weaning and at >1 year old.

### 2.2 Sample collection from wild unknown-age wood mice

Wild wood mice were captured between May 30^th^ – November 23^rd^, 2023, and May 29^th^ – November 21^st^, 2024 at two woodland sites near Edinburgh, Scotland, United Kingdom: Penicuik Estates (55.8261° N, 3.2467° W) and Bilston Wood (55.8714° N, 3.1470° W). At each site, two 80m x 80m population-level grids were placed at least 50m apart; at this distance, no mice were trapped on multiple grids. Each grid was divided into 64 trapping stations (10m x 10m), each containing two Sherman live traps (H.B. Sherman 2 × 2.5 × 6.5 in. folding traps, Tallahassee, FL, USA) set under trap covers, resulting in 256 traps set each night.

We live-trapped each site every three weeks for three consecutive nights for a total of 9 trapping sessions (3 consecutive trapping nights each). Autoclaved Sherman live traps were set late each afternoon and collected the following morning. Each trap contained bedding and a bait consisting of a mix of seeds, mealworms, and a carrot slice. At first capture, individuals above 12g were given subcutaneous RFID microchips (125kHz, Francis Scientific Instruments, UK). From October 23^rd^ to November 23^rd,^ 2024, larger microchips were used, which were only given to mice ≥ 14g (125kHz, WMD, UK). Individuals below these thresholds were given a unique ear biopsy, and if recaptured at a later date above the threshold mass, they received a microchip.

Upon first capture per trapping session, we measured several variables before releasing mice at their site of capture. *Demographic Variables:* We determined sex (based on the appearance of genitalia) and reproductive status. Males with visible descended and scrotal testes were considered reproductive (Herbreteau et al., 2011). Lactating or pregnant females were considered reproductive. We also determined approximate age categories (juvenile, subadult, adult) based on body mass (juvenile: <10g; subadult: 10-15g, adult >15g) and fur colour, which transitions from grey to brown pelage into adulthood (Herbreteau et al., 2011). *Condition*: We measured flattened body length (mm) and body mass (g; using a Pesola scale), with which we calculated skeletal muscle index (proxy of muscle mass) and residual index (deviation from expected mass: length ratio). *Ectoparasites:* We counted the number of ectoparasites including ticks (found primarily on the ears), fleas, and mites (both found at the base of the tail and lower back). *Tissue Samples*: Following ethical guidelines, we also collected a small blood sample from the cheek or tail, which was spun to isolate sera. Every approx. 42 days, we collected a 2mm ear biopsy. Ear biopsies/whole blood and sera were stored immediately at -20 and -80°C, respectively.

Faecal samples were collected from traps within 24-48 hours of capture. First, faecal pellets were weighed and stored in formalin (4% formaldehyde) at 4°C. Approx. 98% of samples were collected by 6pm on the day of capture, i.e., within ≤ 24 hours, with exceptions on heavy trapping days, when a limited number were stored overnight at 4°C and collected the next morning (≤ 48 hours).

#### 2.2.1 Experimental Manipulations

To test for environmental impacts on epigenetic age, we used a two-way factorial experimental design to manipulate both food availability and GI nematode burdens. At each of our two sites, one population-level grid was food-supplemented, and one was given no additional food, resulting in two food-supplemented and two control grids across the study. On our two food supplemented grids, we distributed 10kg of TransBreed^®^ or Safe^®^ 199 high quality nutrient veterinary feeds (protein 18.5%, fat 9.5%, starch 36.8%, high micronutrients, Sweeny, Clerc, et al., 2021) evenly over the grid once a week, starting one to two weeks before the first trapping session. In 2024, each grid experienced the opposite food treatment received in 2023 to account for grid effects. Earlier studies in this system indicate that mice on supplemented grids display better body condition, stronger immune responses, lower parasite loads, and altered parasite communities (Erazo et al., 2025; Sweeny, Clerc, et al., 2021). While individual food consumption cannot be measured, this suggests that supplementation affects physiological traits.

Within each grid, parasite treatment was conducted at the individual level. Wood mice were randomly assigned at first capture to receive either anthelminthic drug treatment or a water control (balanced across sex and time per grid). Anthelminthic-treated individuals received a combination of 100mg/kg Pyrantel and 9.4mg/kg Ivermectin using a sterilised oral gavage. Controls received an equal body size-adjusted amount of tap water. Ivermectin and Pyrantel are common anthelmintic drugs that reduce gastrointestinal nematodes (Pedersen & Fenton, 2015); this combination of drug treatments reduces *H. polygyrus* burdens by >99% in wild wood mice for ∼16 days (Clerc et al., 2019; Sweeny, Albery, et al., 2021). Only individuals large enough to receive RFID tags received treatment, which was repeatedly given at all subsequent recaptures, with a minimum time between doses of 21 days. This design results in four treatments: mice with territories on control or food-supplemented grids who are either given water or anthelminthic treatments every 21+ days.

#### 2.2.2 Sample Selection

Between 2023 and 2024, 461 unique wood mice were captured across our four grids and treated with either water or anthelminthic drugs (see **Table S1** for sample sizes). From those individuals, we generated DNAm from 273 ear biopsies of 163 unique mice. We selected 96 unique mice for whom we had two sampling time points of ear biopsies (typically ≥42 days apart), plus an additional 60 singly captured mice that improved coverage of all age classes and population dynamics, including suspected overwintering adults and singly caught juveniles. In addition, there were 7 individuals who were treated in 2023 and successfully overwintered to be recaptured in 2024.

These individuals each had 2 - 4 DNAm sampling timepoints (total = 21 samples in the dataset). DNAm data from wild individuals was generated on batches (1) and (2) described above, where most 2023 samples were run on batch (1) and 2024 samples (plus additional 2023 samples) were run on batch (2).

### 2.3 Laboratory Methods

#### 2.3.1 Faecal Egg Counts

Faecal samples were analysed using a modified cuvette method (adapted from Hayward et al., 2019). First, the samples were homogenised and then spun down (1500 rpm for 2 minutes) in thin-walled polypropylene tubes, from which the supernatant was aspirated off. The remaining compact faecal matter was thoroughly mixed with a saturated salt solution and spun down again (1500 rpm for 2 minutes). The thin-walled tubes were clamped just below the meniscus using medical forceps, and this top layer of liquid containing GI parasite eggs was poured into a fecalizer.

We allowed oocysts to float onto a coverslip placed on top of the fecalizer for 20 minutes, at which time it was placed onto a slide for analysis. *H. polygyrus e*ggs were counted at 10x magnification. Egg counts were adjusted by the weight of the faecal sample to generate an ‘eggs per gram’ of faeces (EPG). EPG was quantified for all capture samples (n = 1176, n = 1 – 20 samples per individual).

#### 2.3.2 PCR Parasite Presence/Absence

We extracted DNA from whole blood using the automated Kingfisher Flex System (Thermo Fisher Scientific) with MagMAX™-96 DNA Multi-Sample kit (Invitrogen). We detected the presence of three parasites using nested or semi-nested PCRs: wood mouse herpes virus (WMHV), Trypanosomes, and Bartonella *spp*. (see below for details). All 96-well plates included positive (known presence) and negative (water) control wells and were run on a thermal cycler (MJ research, MJ-220). 5⎧l of PCR products was run on 2% agarose gels stained with SYBR Safe.

*WMHV*: WMHV detection followed methods in Knowles et al. (2012), which utilized ILK+ 5′-ATAAACAACAGCTGGCCATCAA-3’ and KG1+ 5′- CTGACCAGATCCACCCCTTT-3’ primers in the first round and TGV+ 5′- TGTAATTCTGTCTATGGCTTCACAGGAGT-3’ and IGY+ 5′- AAGAGAATCTGTGTCTCCATAAAT-3’ primers in the second. Reactions for both rounds consisted of 13.56μl Nuclease-Free Water (Qiagen, 129114), 5.40μl 5x Taq buffer (Meridian Bioscience, BIO-37111), 3.24μl MgCl, 0.54μl dNTPs (Promega, U1202), 1.08μl of forward/reverse primers, and 0.10μl Taq Polymerase (Qiagen, 201203). 25μl of this mix was combined with 2μl of extracted DNA. The thermal cycle was as follows: 25 or 35 cycles (round 1 vs round 2) of (95.0 °C for 3min, 95.0 °C for 20s, 61.0 °C for 30s, and 72.0 °C for 30s), followed by 72.0 °C for 10min.

*Trypanosomes*: We detected the presence of Trypanosomes using a nested PCR that targets the 530bp section of the 18S rRNA gene (Noyes et al., 1999). First round primers included TRY927F 5’-GAAACAAGAAACACGGGAG and TRY927R 5’- CTACTGGGCAGCTTGGA, and second round primers included SSU561F 5’- TGGGATAACAAAGGAGCA and SSU561R 5’-CTGAGACTGTAACCTCAAAGC.

Each reaction had a total volume of 27μl containing 8.25 μl nuclease-free water, 13.5μl MyTaq Red DNA Polymerase (Meridian Bioscience), 2.16μl MgCl (25mM), 0.54μl each of forward and reverse primers, and 2μl genomic DNA. The thermal cycle was as follows: 94°C for 3min followed by 11 cycles of (94°C for 30s, 60°C for 1min (decreasing by 0.5°C each cycle until 55°C), and 72°C for 90s), followed by 9 or 22 (round 1 vs round 2) cycles of (94°C for 30s, 55°C for 60s, and 72°C for 90s), followed by 72°C for 10min.

*Bartonella*: We used a semi-nested PCR that targets the 16S-23S intergenic spacer region to detect Bartonella *spp*. (Telfer et al., 2005). First round primers were BigF (5′-TTGATAAGCGTGAGGTC-3′) and BogR (5′-TGCAAAGCAGGTGCTCTCCCA-3′), and second round primers were BigF and BigR (5′-TCCCAGCTGAGCTACG-3’). The mix for the first round consisted of 8.80μl Nuclease-Free Water (Qiagen, 129114), 13.5μl MyTaq Red Mix (Meridian Bioscience, BIO25044), 2.16μl MgCl2 (Promega, A3511), and 0.27μl of BigF and BogR primers. For each sample, 25μl of this mix was combined with 2μl of extracted DNA, and was run under the following thermal cycle: 96.0 °C for 3min, 13 cycles of 96.0 °C for 10s, 61.0 °C for 10s (decreased by 0.5 °C every cycle), and 72.0 °C for 50s, followed by 8 cycles of 96.0 °C for 10s, 55.0 °C for 10s, 72.0 °C for 50s. The mix for the second round consisted of 9.80μl Nuclease-Free Water, 13.5μl MyTaq Red Mix, 2.16μl MgCl2, and 0.27μl of BigF and BigR primers. 26μl of this mix was combined with 1μl of DNA product from the first round. Round 2 conditions were as follows: 96°C for 3min, followed by 13 cycles of (96°C for 10s, 61°C for 10s (decreased by 0.5°C every cycle), and 72°C for 50s), followed by 22 cycles of (96°C for 10s, 55°C for 10s, and 72°C for 50s). Bartonella product size varies by species – any bands were considered positive for Bartonella.

#### 2.3.3 DNA Extraction & Methylation

DNA was extracted using the Qiagen Blood and Tissue kit, and then bisulfite converted using the Zymo EZ-96 DNA Methylation-Gold (D5007) or EZ-96 DNA Methylation Kits (D5004). At least 300ng of genomic DNA was used for each sample. Methylation state was determined using the custom HorvathMammalMethylChip40 Illumina methylation array (Arneson et al., 2022) which contains > 37,000 probes in genomic regions thought to be highly conserved across mammalian species.

Normalization of raw methylation data was done using the minfi pipeline to generate beta values for each CpG site across individuals (Fortin et al., 2017). In brief, poor-quality samples and probes were filtered based on detection p-values (> 0.01), removing samples with >20% failed probes and probes with >10% failed samples (1^st^batch: 4937 probes; 2nd: 5539 probes; 3rd: 4613 probes); in total, one sample and 5639 unique probes. Signal intensities were normalized using Noob normalization, and only CpGs passing quality control across all three batches were retained. Batch effects were then corrected using the *ComBat* algorithm on logit-transformed (M-value) data, accounting for age and sex as covariates. Beta values are calculated as the ratio of fluorescence of each methylated probe to the total overall probe intensity (Du et al., 2010). Beta values indicate the degree of methylation at each CpG site, ranging from 0 to 1, in which a value of 0 indicates zero gene copies methylated.

Only probes that mapped onto the wood mouse genome (88.62%; n = 33,220 probes; Accession: PRJEB56948, Knowles et al., 2023) and had functional annotations were included in the final dataset, i.e., 27727 probes out of the >37k on the mammalian methyl array. To do so, we aligned the Mammalian Methylation Array probes to the reference genome using the QUASR package (Gaidatzis et al., 2015), and then annotated CpGs using the ChIPseeker package (Yu et al., 2015).

### 2.4 Statistical Analyses

Statistical analyses were done using R Statistical Software (R version 4.5.1; R Core Team, 2025) and ‘lme4’ package for mixed-effects models (Bates et al., 2015).

#### 2.4.1 Building an epigenetic clock using captive wood mice

Our DNAm dataset of 129 captive wood mouse samples was semi-randomly split into two sets for training (n = 74, 168.7 ± 14.9 (range: 25–383), 38 males, 36 females) and testing (n = 55, 192.3 ± 17.8 (range: 25–383), 27 males, 28 females) the epigenetic clock. Repeatedly sampled mice with two timepoints (from batch #2) were randomly split so that only one sample was included in the training dataset.

We built our epigenetic clock using an elastic net penalized regression model on beta values from all CpG sites that passed quality control (glmnet package, Friedman et al., 2021). We set the elastic net mixing parameter alpha to 0.1, which yielded the lowest median absolute error (MdAE) when compared to alpha values of 0.2 to 1 (tested at increments of 0.1). Model tuning was performed using the cv.glmnet function, which performs leave-one-out cross-validation (LOOCV) to select the optimal regularization parameter (λ). In LOOCV, n models are fit (where n is the number of training samples), each time leaving one sample out, training on the remaining samples, and evaluating prediction error on the held-out sample. Here, λ controls how strongly CpG coefficients are shrunk, and thus the complexity of the model. The final clock is refitted on the full training dataset using the selected λ via glmnet, producing a final set of CpGs and coefficients that define the clock. The accuracy of our epigenetic clock was assessed using Pearson correlation (r) and median absolute error (MdAE) between chronological and predicted epigenetic age.

We also evaluated clock performance by applying it to our test dataset (n = 55). A subset of these samples was derived from individuals also included in the training dataset, but sampled ∼11 months earlier or later, and are therefore not fully independent. To address this, we repeated the analysis using only completely independent individuals (n = 14). Despite the reduced sample size, MdAE and correlation estimates were comparable to those obtained using the full test dataset (**Figure S1**). Finally, we compared the performance of the wood mouse clock with that of the pan-mammalian, multi-tissue epigenetic clock (**Figure S1**). We exported the three pan-mammalian clock coefficients from Lu et al. (2023).

An epigenome-wide association study (EWAS) was performed using the R package *limma* (empirical Bayes moderated t-statistics, Ritchie et al., 2015) to identify CpG sites at which DNAm is associated with chronological age. High genomic inflation was observed (λ = 11.7); therefore, we used the bacon package (van Iterson et al., 2017), which lowered genomic inflation to λ = 1.5. P-values were then FDR-adjusted for multiple testing. CpG sites that were significantly correlated with chronological age (FDR-adjusted p < 0.05) were analyzed for functional enrichment using the Genomic Region of Enrichment Annotation Tool (GREAT; McLean et al., 2010) with a human Hg19 background.

#### 2.4.2 Using epigenetic age as a proxy for chronological age in wild wood mice

We first applied our wood mouse epigenetic clock to estimate chronological age. To evaluate its performance, we fit a linear mixed-effects model testing whether epigenetic age covaries with predictors that have well-established relationships with chronological age. If the expected associations are observed, this would support the validity of applying a captive-derived clock to wild populations with unknown-age individuals. Specifically, we expect epigenetic age to be predicted by:

*Timepoint*: Chronological age will be greater at timepoint 2 vs timepoint 1.

*Julian Date*: Previous work using weight, body length, and pelage (Baker, 1946; Brown, 1953; King, 1950) shows a spring population of overwintering adults followed by the emergence of younger cohorts peaking in later summer. We therefore expect epigenetic age to decrease with date.

*Body Mass*: Older adults should be heavier than younger juveniles.

*Reproductive Activity*: Older adults will more likely be reproductively active.

*Sex*: Given males’ typically shorter lifespans (Lemaître et al., 2020), we expect older age classes to be female-biased.

We also included environmental factors with less clear predictions that may influence the age structure of the population via demographic processes, including *study site, year sampled, and supplemental feeding treatment*. Although anthelmintic treatment was randomly assigned and is not expected to influence chronological age, we included it to account for any effects on biological ageing aspects of this clock. Our dataset included 273 samples from 163 unique mice, and so we included individual ID as a random effect to account for repeated measures.

We also predict that older mice are more likely to be infected with parasites. At each capture, we collected data on seven parasites, defined broadly to include: three ectoparasites (number of ticks, mites, and fleas), three bloodborne microparasites/pathogens (WMHV, *bartonella*, and *trypanosoma*), and faecal egg burden of the GI nematode *H. polygyrus*. We tested whether parasite burdens were associated with epigenetic age in two ways. First, using a generalized linear model (GLM) with a Poisson distribution and log link function, we predicted the number of parasite species detected on each capture as a function of supplemental treatment, epigenetic age, Julian date, sex, reproductive status, and year. Second, we used a multivariate analysis of variance (MANOVA) to predict a composite parasite burden across all seven parasites with the same predictors, followed by post-hoc univariate ANOVAs to identify parasite-specific effects. Analyses included only data collected before each mouse’s first anthelminthic or control treatment.

#### 2.4.3 Assessing drivers of variation in rates of epigenetic ageing

While our wood mouse epigenetic clock can be largely interpreted as an estimate of chronological age, we may gain insight about biological ageing or ageing rates by assessing changes in epigenetic age over time within an individual. Of the 163 unique wild wood mice with predicted epigenetic ages, 101 mice had epigenetic age estimated at two timepoints spaced at least 42 days apart (mean = 78.05 ± 6.30 (SE) days apart; maximum = 344 days; 2023: 38F, 44M; 2024: 6F, 13M). For each mouse, we calculated the difference in epigenetic age and number of days between each mouse’s two timepoints, with which we then calculated the slope of change in epigenetic age as the difference in epigenetic age divided by the duration between samples. A linear model with a Gaussian distribution was used to ask whether the change in epigenetic age matched the time elapsed between each mouse’s two samples, and whether other factors mediated that relationship, including food supplementation, anthelminthic treatment, sex, year, the reproductive status of the mouse at timepoint 1, as well as the duration elapsed. We also included interactions between our treatments, as well as between duration elapsed and each treatment.

## 3. RESULTS

### 3.1 A wood mouse-specific epigenetic clock accurately predicts chronological age

We constructed an epigenetic clock using known-age, laboratory-reared wood mice. We trained the clock on a subset of 74 samples using leave-one-out cross-validation with penalized elastic net regression. This yielded a clock based on 177 CpGs that predicted chronological age with a median absolute error (MdAE) of 5.45 days (r = 0.99, p < 0.0001; **Figure 1A**). Applied to an independent test set of 55 mice that were not used to train the clock, the clock maintained strong performance (MdAE = 14.64 days; r = 0.854, p < 0.0001; **Figure 1B**). In comparison, the universal pan-mammalian clocks (Lu et al., 2023) are less accurate, with some estimates reaching 1000+ days of age (MdAE = 152-175 days; r = 0.86-0.88; **Figures 1C, S2**).

**Figure 1.**
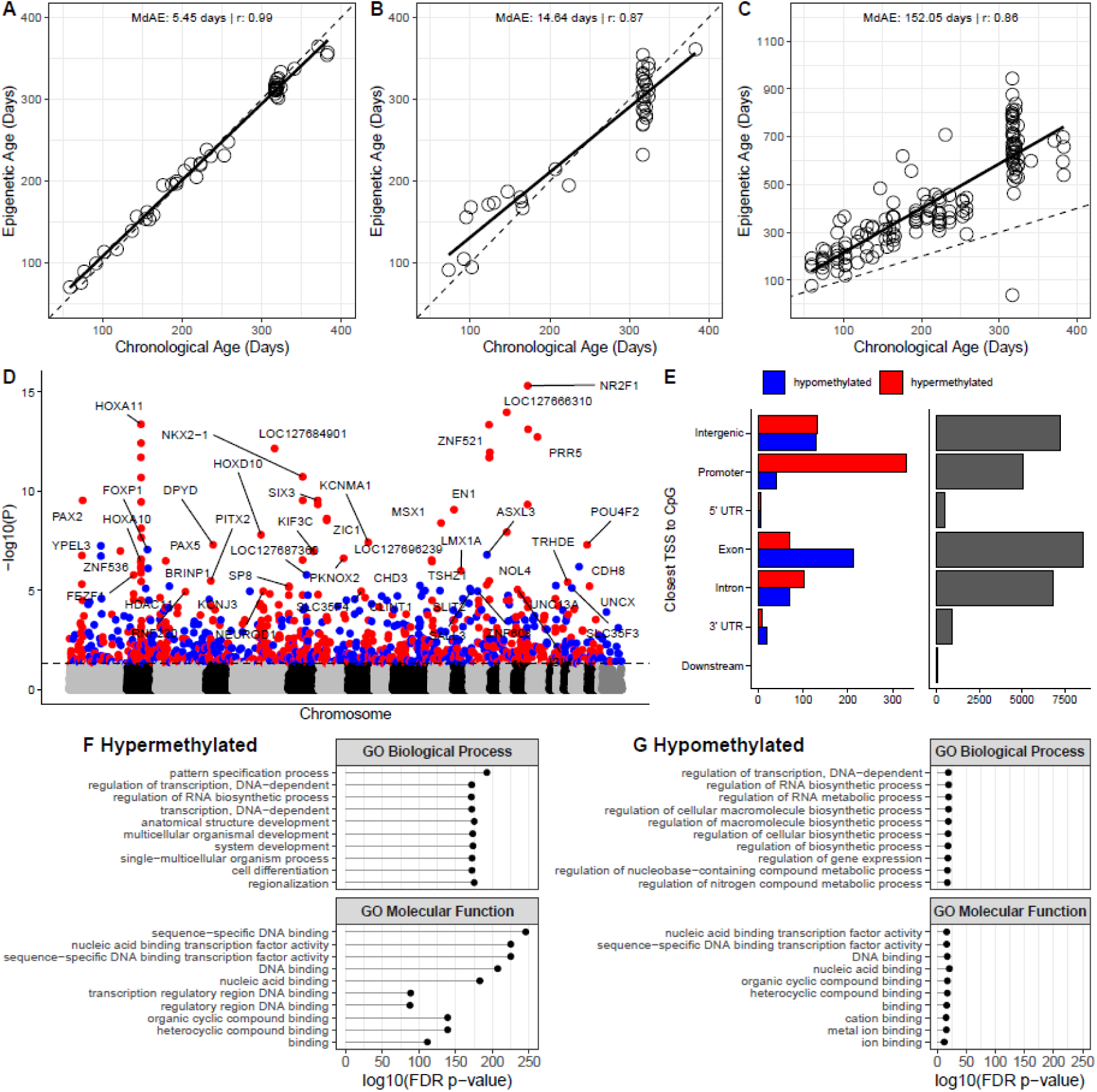
Development of the wood mouse epigenetic clock. (A) n = 74 mice used for clock training; (B) n = 55 mice used for independent testing of the wood mouse clock. (C) Estimated epigenetic age from the Universal Pan-Mammalian Clock #2 (relative chronological age). The diagonal dashed lines depict y = x. (D) Manhattan plot of CpG sites significantly correlated with chronological age (FDR-adjusted p < 0.05 following adjustments for genomic inflation). CpGs are coloured by direction of association (red = hypermethylated; blue = hypomethylated). Only the top 50 unique CpGs in each direction are labelled with their associated gene name. (E) Location of significant age-related CpGs relative to the adjacent transcriptional start site. Gray bars show CpG locations of all CpG sites on the mammalian methyl array that map to the wood mouse genome. (F, G) The top 10 terms per database are shown for significantly (F) hyper- and (G) hypomethylated groups of CpG sites (FDR-adjusted p < 0.05).

An epigenome-wide association study (EWAS) across all quality-controlled CpGs identified 1,127 age-associated sites (FDR < 0.05), comprising 651 hypermethylated and 476 hypomethylated CpGs (**Figure 1D**). Age-associated hypermethylated CpGs were often found in promoter regions (**Figure 1E**). GREAT analysis (Genomic Regions Enrichment of Annotations Tool) indicated that hypermethylated CpGs were associated with gene ontology terms related to system development, pattern specification, and differentiation, as well as DNA binding and transcription factor activity. The top five hypermethylated genes were *NR2F1*, *LOC127666310*, *HOXA11*, *ZNF521*, and *PRR5*. These sites also showed enrichment near polycomb repressive complex 2 (PRC2) targets (e.g., SUZ12, EED, H3K27me3), indicating epigenetic regulation of developmental genes. In contrast, hypomethylated CpGs were enriched for gene sets involved in transcription, biosynthesis, and gene expression (**Figure 1F**). The top five hypomethylated genes were *YPEL3*, *FOXP1*, *ASXL3*, *CDH8*, and *LOC127687360*.

### 3.2 DNAm captures chronological age in the wild

We applied the wood mouse epigenetic clock to ear biopsies from wild-caught mice to infer chronological age. The clock predicted ages spanning 11.99 to 397.10 days, with an average of 135 days. In addition, small (<12 gram), grey juveniles – expected to be ∼20–50 days old – showed clock-based ages of 43.0 ± 4.15 days (**Figure 2A**). For comparison, age estimates using the Pan-Mammalian clocks (Lu et al., 2023) averaged ∼320 days with ranges of ∼35 to 1400 days-old (**Figure S3**). In a mixed-effects model (273 samples, 163 mice), we also tested whether epigenetic age tracks well-established correlates of chronological age. As predicted, on average epigenetic age was higher at individuals’ second sampling timepoints (β = 60.99 ± 8.60, F_222.53_ = 49.75, p < 0.001), declined over the breeding season – consistent with recruitment of younger cohorts (β = -119.58 ± 32.08, F_261.62_ = 13.72, p < 0.001), and increased with body mass (β = 7.37 ± 1.09, F_261.15_ = 45.42, p < 0.001, **Figure 2B, 2C**). Epigenetic age did not differ by sex or current reproductive status, nor between experimental groups, as expected under randomised treatments (p ≥ 0.15, see treatment details below). Age structure also varied spatially and temporally (site: F_153.38_ = 5.82, p = 0.018; year: F_213.23_ = 10.20, p = 0.002).

**Figure 2.**
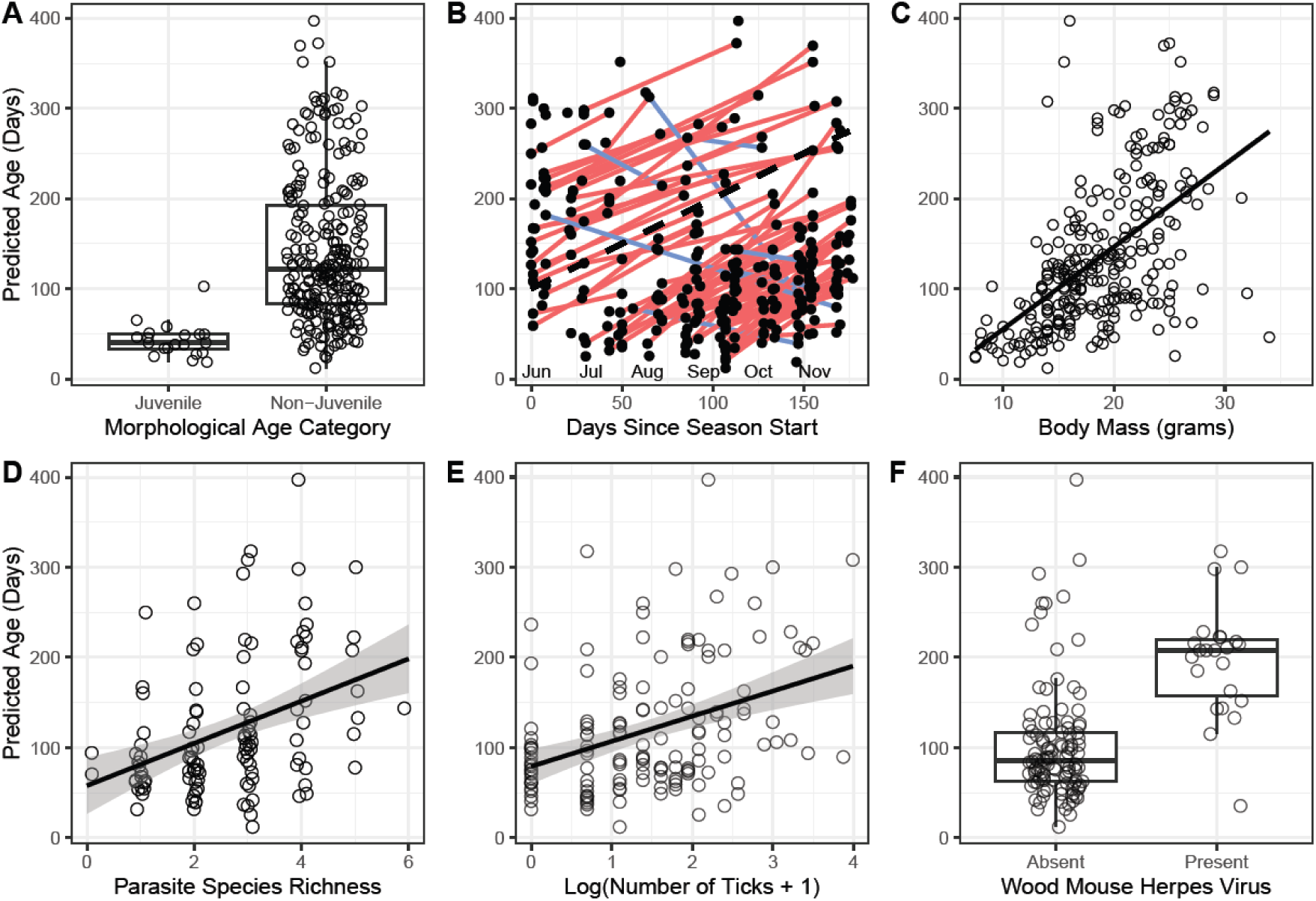
Significant correlates of predicted age in wild wood mice. **(A)** Small (<12 g), grey juveniles - expected to be ∼20-50 days old - were predicted to be younger than non-juveniles. **(B)** Predicted age generally increased with sampling date (lines connect repeated samples for the same individual; red/blue lines signify epigenetic age increasing/decreasing, respectively) and **(C)** body mass. **(D)** Parasite species richness (number of unique parasite species per individual) also increased with predicted age. Post-hoc analyses revealed that older-predicted mice harboured **(E)** more ticks and **(F)** had a higher probability of wood mouse herpes virus infection.

### 3.3 Estimates of chronological age capture age-related patterns of parasitism

As an independent validation of the clock’s functionality in wild mice, we examined whether baseline parasite burden varies with epigenetic age. Epigenetic age was positively associated with parasite diversity (β = 0.00195 ± 0.00079, *z* = 2.47, *p* = 0.0135; **Figure 2D; Table S2**). A MANOVA predicting a composite metric of parasite burden from all seven parasites also showed a strong multivariate association with epigenetic age (Pillai’s trace = 0.373, F_₇,₁₀₂_ = 8.66, *p* < 0.0001; **Table S3**). Post hoc univariate tests revealed that older mice had higher tick burdens (*F* = 18.89, *p*_adj_ = 0.0006) and a greater likelihood of WMHV infection (*F* = 38.92, *p*_adj_ < 0.0001; **Figure 2E, 2F; Table S4**), with other parasite burdens not strongly related to age.

### 3.4 Epigenetic ageing decelerates in response to food supplementation and parasite removal in the wild

We also conducted a two-factor field experiment across two forest sites to test whether environmental factors influence epigenetic ageing. We found that food supplementation increased mouse abundance and both treatments reduced worm egg burdens, but there were no effects on reproductive or body condition (see Supplementary Information for detailed analyses, **Tables S5-6**, **Figures S4-S5**).

We conducted a longitudinal analysis focused on 101 experimental wild mice (a subset of the individuals described above) who were sampled twice (typically ≥42 days apart; mean 78.1 ± 6.3 SE days; maximum 344 days). For each day lived, mice should age by one epigenetic day. Across all individuals, the slope of epigenetic change averaged 0.89 ± 0.07 epigenetic per chronological day (range −2.65 to 2.80), close to the expected slope of 1 (**Figure 3A**). Epigenetic ageing increased with the time elapsed between samples (β = 0.90 ± 0.15, F = 34.88, p < 0.0001) in the vast majority of mice. We also observed a significant interaction between elapsed time and our combinations of food supplementation and anthelmintic treatment (β = -0.41 ± 0.17, F = 5.57, p = 0.001, **Figure 3B**). Specifically, post hoc analyses show that it is the combination of both treatments (supplemental food and parasite removal) that significantly slows the rate of epigenetic ageing (**Table S7**). Epigenetic ageing was not associated with year, sex, or initial reproductive status (p ≥ 0.25; **Table S8**).

**Figure 3.**
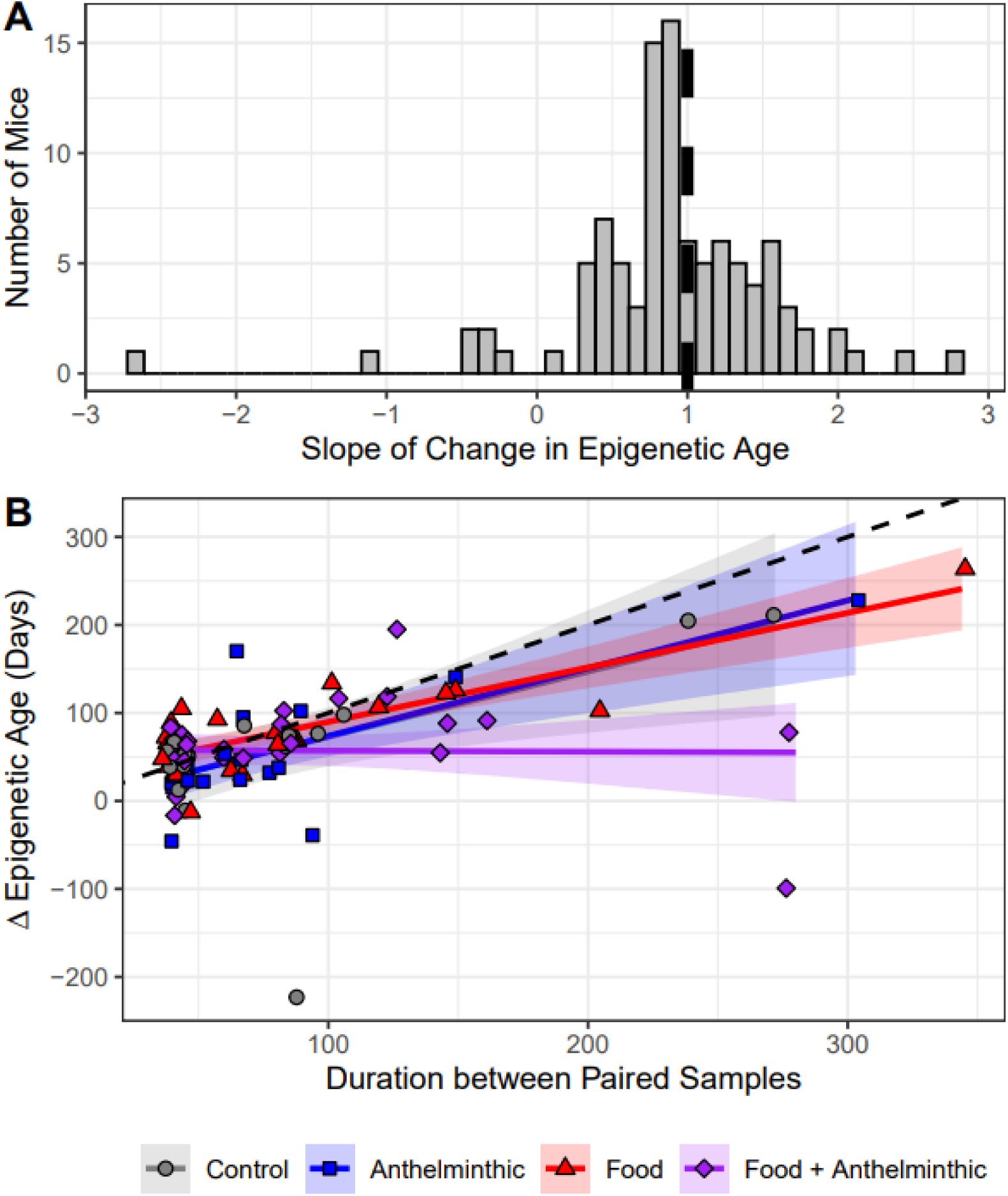
The change in epigenetic age over time in wild wood mice. Analyses focus on the 101 mice in the dataset with ≥2 epigenetic age measurements. **(A)** Histogram of individual slopes, indicating how closely changes in epigenetic age correspond to the number of days elapsed. **(B)** Effects of food supplementation and anthelminthic treatments on the change in epigenetic age over time. Points are jittered for clarity. The black dashed line represents the expected slope of 1, where epigenetic ageing matches chronological time elapsed.

## 4. DISCUSSION

Despite the potential of short-lived, experimentally tractable wild systems for ageing research, deploying epigenetic clocks in the wild remains challenging, particularly when chronological age is unknown. Here, we present one of the first examples of a laboratory-trained, species-specific epigenetic clock successfully deployed in the wild, using the wood mouse as a model. In captive mice, the clock predicted age with high accuracy (MdAE = 14.64 days). Age-associated CpGs were predominantly hypermethylated, located in promoter regions, and linked to developmental processes. Critically, the clock performs robustly in wild populations, as evidenced by multiple independent validations: epigenetic ages (1) fell within expected age ranges, particularly for juveniles whose chronological age is most accurately estimable, (2) reflect population-level demographic patterns across the breeding season, and (3) increase over time at an average slope of ∼1 unit/day. Furthermore, mice receiving both food supplementation and anthelmintic treatment – reducing two key environmental stressors – exhibited slower epigenetic ageing, indicating that the clock captures aspects of biological, not just chronological, age.

Species-specific epigenetic clocks generally provide more accurate age estimates (Parsons et al., 2023). In our study, chronological age in laboratory-reared wood mice was better predicted by the species-specific clock than by the pan-mammalian clock (MdAE of 15 versus 175 days), which overestimated chronological age in some cases by 3x, up to ∼1000 days old, suggesting that the application of pan-mammalian clocks to new datasets may be nontrivial, potentially requiring complex data conversions. Regardless, with the wood mouse clock, a 15-day error is only 0.68% of a 6-year maximum lifespan (as estimated by AnAge, De Magalhaes & Costa, 2009), 4% of a 1-year maximum lifespan of wild mice, or 17% of a shorter, more typical 3-month lifespan in the wild. Although an error of ≤20% may seem high, this represents a substantial improvement over conventional morphology-based age estimates, with which a 60-day-old and 360-day-old adult are effectively indistinguishable. 26 of the 177 CpGs in our clock are also included in at least one of the pan-mammalian clocks (Lu et al., 2023). The five genes associated with CpGs in all three clocks include *YPEL3*, *OTP*, *ZIC1*, *TNRC6A*, and *NKX2-3*. While this likely underestimates the overlap, these comparisons can start to shed light on conserved mechanisms of ageing. It is also worth noting that our use of the mammalian methylation array (Arneson et al., 2022), which only assesses 37k CpG sites, may exclude key species-specific age-associated sites not represented on the array.

Analysis of age-associated hyper- and hypomethylated CpGs can reveal key pathways contributing to senescence. Among the top hypermethylated CpGs were *NR2F1*, *HOXA11*, *ZNF521*, and *PRR5*. Increased DNAm of *NR2F1* and *ZNF521* may suppress developmental and stemness genes, consistent with age-related decline in regenerative capacity (Bond et al., 2004; Bonzano et al., 2023). Epigenetic silencing of HOX genes (e.g., *HOXA11*) in adulthood also aligns with these expectations (Pearson et al., 2005), and notably, *PRR5*, a component of mTOR signaling, may contribute to metabolic and insulin signaling decline with age (Pearce et al., 2007). Critically, many hypermethylated CpGs occur near genes regulated by the polycomb repressive complex 2 (i.e., PRC2, Bracken et al., 2006), which is tightly linked to development. Indeed, epigenetic clocks frequently include PRC2 targets (Dozmorov, 2015; Horvath, 2013; Robeck et al., 2023; Zoller et al., 2024). In contrast, several of the most hypomethylated CpGs mapped to *YPEL3*, *FOXP1*, and *ASXL3*. *YPEL3* acts downstream of p53 to induce cellular senescence and may function as a tumour suppressor that becomes more active with age (Kelley et al., 2010). ASXL3 plays a key role in histone modification and may influence age-related PRC2 activity (Abdel-Wahab et al., 2012; Shukla et al., 2017), while *FOXP1* regulates developmental and immune pathways, though its expression typically decreases with age (Benisch et al., 2012; Li et al., 2017).

Our wood mouse epigenetic clock appears to estimate chronological age remarkably well in wild individuals, with predicted ages falling within the expected range – under one year old and averaging 135 days (4 – 5 months). This aligns with expectations for short-lived wild systems, where extrinsic mortality can be quite high (Collins & Kays, 2014) and subsequently, a large proportion dies well before their maximum lifespan is reached. Notably, the clock places juveniles within their expected age range (20–50 days), showing high accuracy among the youngest individuals, whose age can be confidently estimated from their body size and distinctive colour patterns. The upper limit of adult age estimates also match expectations, but we do not currently know how precise these estimates are. Greater uncertainty is likely in older age classes if epigenetic ageing rates fluctuate from development into adulthood (Snir et al., 2019). In addition, underestimation may occur if wild mice age faster than the colony-reared individuals used to build the clock, as suggested by work in mice (Hanski et al., 2024). Even so, the clock provides far more precise estimates than those gained from morphology.

Even if the clock does not perfectly estimate absolute chronological age, relative age estimates remain highly informative for understanding population structure. In wild wood mice, the first individuals to appear in May-June are typically old overwintering survivors from the previous breeding season, followed by a breeding peak in August–September and a surge of juveniles captured in October. These dynamics are clearly reflected in our epigenetic age estimates across the breeding season: although epigenetic age increases within individuals over time, population-level averages decline as the younger cohort emerges. Thus, epigenetic clocks can transform demographic monitoring by revealing shifts in age structure and enabling the reconstruction of a complete demographic matrix from a single cross-sectional sampling event (Eichenberger et al., 2026; Zhao et al., 2018). Such approaches can, in turn, inform key parameters of population dynamics, including growth rate – an essential metric for IUCN conservation assessments (Mace et al., 2008).

Moreover, parasite richness, tick burden, and wood mouse herpes virus (WMHV) infection probability were positively associated with epigenetic age, suggesting that older mice face greater exposure and susceptibility to infection. In the wild, older individuals often harbour more parasites due to cumulative exposure and life-history differences in habitat use, movement, and social interactions that influence contact rates (Albery et al., 2023). Age-related declines in immune function, called immunosenescence, further reduce the ability to control infections and are well documented across taxa (Humphreys & Grencis, 2002; A. Peters et al., 2019).

Consistent with this, WMHV behaves as a chronic, age-associated infection in wood mice (Erazo et al., 2021). In addition, parasites may also accelerate epigenetic ageing (Bai et al., 2026; Durso et al., 2022; Gupta & Vadde, 2026) via energetic costs, tissue damage, and modulation of host physiology, potentially altering DNAm patterns (McNew et al., 2021; Sagonas et al., 2020). While we cannot determine causality or the relative contributions of these mechanisms, our results confirm the expected positive association between parasitism and epigenetic age. Notably, the epigenetic clock enables us to capture these age-parasite relationships with unprecedented resolution, facilitating refined tests of age–infection dynamics in the wild.

Few studies have applied epigenetic clocks longitudinally in wild populations (Anderson et al., 2021; De Paoli Iseppi et al., 2019; Hanski et al., 2024; Pinho et al., 2022; Sullivan et al., 2022). Here, epigenetic age increased at an average rate of 0.89 ± 0.07 units per day – close to the expected 1:1 relationship with chronological time. Only 7% of mice showed decreases in epigenetic age, similar to the small fraction of apparent “rejuvenators” reported in other systems (De Paoli Iseppi et al., 2019; e.g., 1/7 birds and 2/14 marmots, both older individuals, Pinho et al., 2022). Most declines were modest (slope ∼ -0.40) and may reflect known plasticity in epigenetic ageing, such as rank-related decreases observed in baboons (Anderson et al., 2021). Alternatively, such patterns may reflect transient shifts in tissue composition (e.g., immune cell infiltration or wound healing in ear biopsies) or short-term physiological or environmental effects (Jaffe & Irizarry, 2014; Pastar et al., 2021). Tissue type may also influence epigenetic ageing rates; ear biopsies, while heterogeneous, may be less responsive in DNAm dynamics than blood, which is highly labile due to shifts in immune cell composition and rapid cell turnover.

Encouragingly, epigenetic age estimates are strongly correlated across tissues, supporting the robustness of our ear-based clock (**Figure S5**). A few mice did show much larger declines (e.g., 100 units lost over 280 days, 223 units lost over 84 days). The causes of these extreme cases remain unclear – they may reflect environmental effects or technical artifacts – but were retained in analyses for completeness.

One logical prediction is that wild mice will epigenetically age faster than laboratory mice due to the greater ecological and physiological challenges of natural environments (Zipple et al., 2025), as shown in *Mus* spp. (Hanski et al., 2024).

However, about 50% of our slopes fell between 0 and 1, suggesting that wild mice aged more slowly than predicted by a lab-derived clock. This pattern could reflect clock biases, such as underestimation in older individuals due to nonlinear DNA methylation changes (Dufek et al., 2024) or regression to the mean (Barnett et al., 2005). Biological factors may also contribute: overwinter torpor (see below), smaller body size, and differences in growth and metabolism could slow epigenetic ageing in wild mice, while lab mice often experience nutrient-rich but uniform diets, reduced activity, and faster growth, all of which can accelerate epigenetic ageing (Galkin et al., 2023; Quach et al., 2017). Despite potential underestimation of epigenetic ageing rates, the clock still provides meaningful insights into ageing biology within wild populations.

Our results suggest that experimental manipulation of food availability and parasite burdens can slow the rate of epigenetic ageing in wild mice. In particular, we found evidence for an additive effect, where individuals receiving both supplemental food and anthelmintic treatment experienced the least epigenetic ageing. This aligns with growing evidence that environmental conditions influence epigenetic ageing dynamics across species – for example, related to social rank (Anderson et al., 2021), hibernation (Pinho et al., 2022; Sullivan et al., 2022), and sibling competition (Haller et al., 2025). We also detected a duration-by-treatment interaction, indicating that treatment effects became apparent only across longer intervals between sampling, suggesting that benefits accumulate over time. Because relatively few individuals were recaptured across long durations, future studies should continue to evaluate these long-term patterns.

We predicted that alleviating stress from food scarcity and parasite infection would slow epigenetic ageing. Supplemental feeding is well known to increase reproduction and survival, though it can also elevate pathogen transmission risk (Boutin, 1990; Murray et al., 2016). In our study, supplementation increased mouse density, but whether this reflected elevated reproduction or immigration remains unclear.

Because food was distributed evenly throughout the population rather than concentrated at feeding stations, wood mice likely benefited nutritionally without elevated contact rates or infection risk. Gastrointestinal helminths, common nematodes that infect wild mice, can impair host condition, reproduction, and survival through energetic costs, tissue damage, and immune modulation (Maizels & McSorley, 2016; Schwanz, 2008; Scott, 2023; Shanebeck et al., 2022). Together, our treatments may reduce energetic constraints and physiological stress, allowing greater investment in maintenance. Whether this translated into measurable deceleration of epigenetic ageing likely depended on how additional energy was allocated – for example, toward reproduction versus repair (Ryan et al., 2024). Moreover, variation in anthelmintic treatment frequency and food intake among individuals may dilute treatment effects, as treatments are only effective for ∼21 days and rapid reinfection is common (Clerc et al., 2019).

This pattern of treatment-induced epigenetic age deceleration may be driven by the overwintering survivors recaptured ≥ 200 days after their first sample. During winter, wood mice undergo spontaneous daily torpor, a common physiological state that conserves energy via lower metabolic rates during cold or food-scarce periods (Walton & Andrews, 1981). Hibernation often slows biological ageing (Pinho et al., 2022; Power et al., 2023; Sullivan et al., 2022; Turbill et al., 2012), which could explain the study-wide pattern of epigenetic age deceleration. However, frequent arousals from torpor can elevate oxidative damage and accelerate telomere shortening (Geiser, 2004; Hoelzl et al., 2016). Here, our treatments may have improved body condition, or alternatively, increased winter food caches. In this scenario, less reliance on physiologically stressful torpor-arousal cycles may slow epigenetic ageing while maintaining the energetic stability necessary for overwinter survival. However, our study cannot address epigenetic ageing during winter, which would likely require controlled laboratory experiments with access to mice experiencing torpor.

Our epigenetic clock represents a clear advance over traditional ageing methods, providing far finer resolution of age classes, and in doing so, unlocks this system’s potential to reveal how the environment shapes biological ageing in the wild. Until now, most wildlife epigenetic clock studies have focused on building new clocks for new species - an essential step to developing the methodological foundations of reliable clock construction that nevertheless does not link epigenetic age to biological variation. Our study marks a shift in the role of epigenetic clocks in wildlife research: we show how these tools can illuminate individual variation in epigenetic age, quantify age acceleration in both natural and manipulated contexts, and resolve population structure in systems where chronological age is otherwise inaccessible.

As the field moves toward using clocks to address ecological and evolutionary questions, our work helps define this new phase and demonstrates the transformative potential of epigenetic clocks in wildlife biology.

## Acknowledgements

We thank Jessica Hall and Rowan Bancroft for assistance with sample collection from the wood mouse colony and subsequent lab work. We are also grateful for support in the field and laboratory: in 2023, to Mairéad Corr, Alexandra Vavrik, Thomas Loebel-Messer, Isabel Entwistle, Rhoslyn Howroyd, Anders Erlandson, Cara Duffy, and Nathan Loebel-Messer; and in 2024, to Alexandra Vavrik, Thomas Loebel-Messer, Nathan Loebel-Messer, Beth Wilson, Cara Duffy, and Andreia Batista Pereira. We also thank Cameron Watkins and James Nixon for access to field sites. This research was supported by a NERC grant (NE/X001423/1) to TL, ABP, and SAB and a University of Edinburgh SEED award grant (561430) to TL and ABP.

## Data Availability Statement

Data will be available in the Zenodo Digital Repository.

## Benefit-Sharing Statement

Benefits from this research accrue from the sharing of our data and results on public databases as described above.

## Author Contributions

Sarah E. Wolf, Tom J. Little, Steve Horvath, and Amy B. Pedersen designed the laboratory colony study, and Tom J. Little, Amy B. Pedersen, and Simon A. Babayan designed the study resulting in wild mouse methylation data. Sarah E. Wolf conducted statistical analyses, with consultation from Riccardo E. Marioni. Sarah E. Wolf drafted the manuscript, on which all authors commented and gave final approval for publication.

## REFERENCES

Abdel-Wahab, O., Adli, M., LaFave, L. M., Gao, J., Hricik, T., Shih, A. H., Pandey, S., Patel, J. P., Chung, Y. R., & Koche, R. (2012). ASXL1 mutations promote myeloid transformation through loss of PRC2-mediated gene repression. Cancer Cell, 22(2), 180–193.

Albery, G. F., Sweeny, A. R., & Webber, Q. (2023). How behavioural ageing affects infectious disease. Neuroscience & Biobehavioral Reviews, 155, 105426.

Anderson, J. A., Johnston, R. A., Lea, A. J., Campos, F. A., Voyles, T. N., Akinyi, M. Y., Alberts, S. C., Archie, E. A., & Tung, J. (2021). High social status males experience accelerated epigenetic aging in wild baboons. Elife, 10, e66128.

Arneson, A., Haghani, A., Thompson, M. J., Pellegrini, M., Kwon, S. B., Vu, H., Maciejewski, E., Yao, M., Li, C. Z., & Lu, A. T. (2022). A mammalian methylation array for profiling methylation levels at conserved sequences. Nature Communications, 13(1), 783.

Bai, R., Pan, L., Huang, Y., Jiang, Z., Wang, J., Liu, Y., Huang, C., Zheng, X., Yu, Y., & Li, Q. (2026). Association of accelerated biological aging and frailty with the risk of severe infection: A prospective study in the UK Biobank. The Journal of Frailty & Aging, 15(1), 100118.

Baker, R. H. (1946). A study of rodent populations on Guam, Mariana Islands. Ecological Monographs, 16(4), 393–408.

Barnett, A. G., Van Der Pols, J. C., & Dobson, A. J. (2005). Regression to the mean: What it is and how to deal with it. International Journal of Epidemiology, 34(1), 215–220.

Bates, D., Mächler, M., Bolker, B., & Walker, S. (2015). Fitting linear mixed-effects models using lme4. Journal of Statistical Software, 67, 1–48.

Benisch, P., Schilling, T., Klein-Hitpass, L., Frey, S. P., Seefried, L., Raaijmakers, N., Krug, M., Regensburger, M., Zeck, S., & Schinke, T. (2012). The transcriptional profile of mesenchymal stem cell populations in primary osteoporosis is distinct and shows overexpression of osteogenic inhibitors.

Bentz, A. B., George, E. M., Wolf, S. E., Rusch, D. B., Podicheti, R., Buechlein, A., Nephew, K. P., & Rosvall, K. A. (2021). Experimental competition induces immediate and lasting effects on the neurogenome in free-living female birds. Proceedings of the National Academy of Sciences, 118(13).

Bond, H. M., Mesuraca, M., Carbone, E., Bonelli, P., Agosti, V., Amodio, N., De Rosa, G., Di Nicola, M., Gianni, A. M., & Moore, M. A. (2004). Early hematopoietic zinc finger protein (EHZF), the human homolog to mouse Evi3, is highly expressed in primitive human hematopoietic cells. Blood, 103(6), 2062–2070.

Bonzano, S., Dallorto, E., Molineris, I., Michelon, F., Crisci, I., Gambarotta, G., Neri, F., Oliviero, S., Beckervordersandforth, R., & Lie, D. C. (2023). NR2F1 shapes mitochondria in the mouse brain, providing new insights into Bosch-Boonstra-Schaaf optic atrophy syndrome. Disease Models & Mechanisms, 16(6), dmm049854.

Boutin, S. (1990). Food supplementation experiments with terrestrial vertebrates: Patterns, problems, and the future. Canadian Journal of Zoology, 68(2), 203–220.

Bracken, A. P., Dietrich, N., Pasini, D., Hansen, K. H., & Helin, K. (2006). Genome-wide mapping of Polycomb target genes unravels their roles in cell fate transitions. Genes & Development, 20(9), 1123–1136.

Brown, R. Z. (1953). Social behavior, reproduction, and population changes in the house mouse (Mus musculus L.). Ecological Monographs, 23(3), 218–240.

Clerc, M., Babayan, S. A., Fenton, A., & Pedersen, A. B. (2019). Age affects antibody levels and anthelmintic treatment efficacy in a wild rodent. International Journal for Parasitology: Parasites and Wildlife, 8, 240–247.

Collins, C. R., & Kays, R. W. (2014). Patterns of mortality in a wild population of white-footed mice. Northeastern Naturalist, 21(2), 323–336.

Cristofari, R., Davis, L. R., Bardon, G., Nitta Fernandes, F. A., Figueroa, M. E., Franzenburg, S., Gauthier-Clerc, M., Grande, F., Heidrich, R., Hukkanen, M., Le Maho, Y., Ollikainen, M., Paciello, E., Rampal, P., Stenseth, N. C., Trucchi, E., Zahn, S., Le Bohec, C., & Meyer, B. S. (2026). Lifestyle change accelerates epigenetic ageing in King penguins. Nature Communications. 10.1038/s41467-026-70527-8

De Magalhaes, J., & Costa, and J. (2009). A database of vertebrate longevity records and their relation to other life-history traits. Journal of Evolutionary Biology, 22(8), 1770–1774.

De Paoli-Iseppi, R., Deagle, B. E., Polanowski, A. M., McMahon, C. R., Dickinson, J. L., Hindell, M. A., & Jarman, S. N. (2019). Age estimation in a long-lived seabird (Ardenna tenuirostris) using DNA methylation-based biomarkers. Molecular Ecology Resources, 19(2), 411–425.

Dozmorov, M. G. (2015). Polycomb repressive complex 2 epigenomic signature defines age-associated hypermethylation and gene expression changes. Epigenetics, 10(6), 484–495.

Du, P., Zhang, X., Huang, C.-C., Jafari, N., Kibbe, W. A., Hou, L., & Lin, S. M. (2010). Comparison of Beta-value and M-value methods for quantifying methylation levels by microarray analysis. BMC Bioinformatics, 11, 1–9.

Dufek, G., Katriel, G., Snir, S., & Pellegrini, M. (2024). Exponential dynamics of DNA methylation with age. Journal of Theoretical Biology, 579, 111697.

Durso, D., Silveira-Nunes, G., Coelho, M., Camatta, G., Ventura, L., Nascimento, L., Caixeta, F., Cunha, E., Castelo-Branco, A., & Fonseca, D. M. da. (2022). Living in endemic area for infectious diseases accelerates epigenetic age. Mechanisms of Ageing and Development, 207, 111713.

Eichenberger, F., Carroll, E. L., Garrigue, C., Jarman, S., Steel, D. J., Robbins, J., Rendell, L., & Garland, E. C. (2026). Changes in age-related sexual selection in a humpback whale population recovering from exploitation. Current Biology.

Erazo, D., Pedersen, A. B., Gallagher, K., & Fenton, A. (2021). Who acquires infection from whom? Estimating herpesvirus transmission rates between wild rodent host groups. Epidemics, 35, 100451.

Erazo, D., Sweeny, A. R., Pedersen, A. B., & Fenton, A. (2025). Parasite responses to resource provisioning can be altered by within-host co-infection interactions. Proceedings of the Royal Society B: Biological Sciences, 292(2058).

Esteban, M., García-París, M., & Castanet, J. (1996). Use of bone histology in estimating the age of frogs (Rana perezi) from a warm temperate climate area. Canadian Journal of Zoology, 74(10), 1914–1921.

Fortin, J.-P., Triche Jr, T. J., & Hansen, K. D. (2017). Preprocessing, normalization and integration of the Illumina HumanMethylationEPIC array with minfi. Bioinformatics, 33(4), 558–560.

Friedman, J., Hastie, T., Tibshirani, R., Narasimhan, B., Tay, K., Simon, N., & Qian, J. (2021). Package ‘glmnet.’ CRAN R Repositary, 595.

Gaidatzis, D., Lerch, A., Hahne, F., & Stadler, M. B. (2015). QuasR: quantification and annotation of short reads in R. Bioinformatics, 31(7), 1130–1132.

Galkin, F., Kovalchuk, O., Koldasbayeva, D., Zhavoronkov, A., & Bischof, E. (2023). Stress, diet, exercise: Common environmental factors and their impact on epigenetic age. Ageing Research Reviews, 88, 101956.

Geiser, F. (2004). Metabolic rate and body temperature reduction during hibernation and daily torpor. Annu. Rev. Physiol., 66, 239–274.

Gupta, M. K., & Vadde, R. (2026). DNA methylation-based epigenetic clocks highlight immune-driven aging acceleration in COVID-19 across diverse populations. Biogerontology, 27(1), 12.

Haller, A., Risse, J., Sepers, B., & van Oers, K. (2025). Independent Avian Epigenetic Clocks for Ageing and Development. Molecular Ecology Resources, e14128.

Hanski, E., Joseph, S., Raulo, A., Wanelik, K. M., O’Toole, Á., Knowles, S. C., & Little, T. J. (2024). Epigenetic age estimation of wild mice using faecal samples. Molecular Ecology, 33(8), e17330.

Hayward, A. D., Pilkington, J. G., Wilson, K., McNeilly, T. N., & Watt, K. A. (2019). Reproductive effort influences intra-seasonal variation in parasite-specific antibody responses in wild Soay sheep. Functional Ecology, 33(7), 1307–1320.

Herbreteau, V., Jittapalapong, S., Rerkamnuaychoke, W., Chaval, Y., Cosson, J.-F., & Morand, S. (2011). Protocols for field and laboratory rodent studies. Kasetsart University.

Hoelzl, F., Cornils, J. S., Smith, S., Moodley, Y., & Ruf, T. (2016). Telomere dynamics in free-living edible dormice (Glis glis): The impact of hibernation and food supply. Journal of Experimental Biology, 219(16), 2469–2474.

Horvath, S. (2013). DNA methylation age of human tissues and cell types. Genome Biology, 14, 1–20.

Horvath, S., & Raj, K. (2018). DNA methylation-based biomarkers and the epigenetic clock theory of ageing. Nature Reviews Genetics, 19(6), 371–384.

Humphreys, N. E., & Grencis, R. K. (2002). Effects of ageing on the immunoregulation of parasitic infection. Infection and Immunity, 70(9), 5148–5157.

Jaffe, A. E., & Irizarry, R. A. (2014). Accounting for cellular heterogeneity is critical in epigenome-wide association studies. Genome Biology, 15(2), R31.

Kelley, K. D., Miller, K. R., Todd, A., Kelley, A. R., Tuttle, R., & Berberich, S. J. (2010). YPEL3, a p53-regulated gene that induces cellular senescence. Cancer Research, 70(9), 3566– 3575.

King, O. M. (1950). An ecological study of the Norway rat and the house mouse in a city block in Lawrence, Kansas. Transactions of the Kansas Academy of Science (1903-), 53(4), 500–528.

Knowles, S. C., Fenton, A., & Pedersen, A. B. (2012). Epidemiology and fitness effects of wood mouse herpesvirus in a natural host population. Journal of General Virology, 93(11), 2447–2456.

Knowles, S. C., Raulo, A., of Oxford, U., Lab, W. W. G. A., of Life, W. S. I. T., & Darwin Tree of Life Consortium. (2023). The genome sequence of the wood mouse, Apodemus sylvaticus (Linnaeus, 1758). Wellcome Open Research, 8, 442.

Le Clercq, L., Kotzé, A., Grobler, J. P., & Dalton, D. L. (2023). Biological clocks as age estimation markers in animals: A systematic review and meta-analysis. Biological Reviews, 98(6), 1972–2011.

Lemaître, J.-F., Ronget, V., Tidière, M., Allainé, D., Berger, V., Cohas, A., Colchero, F., Conde, D. A., Garratt, M., & Liker, A. (2020). Sex differences in adult lifespan and aging rates of mortality across wild mammals. Proceedings of the National Academy of Sciences, 117(15), 8546–8553.

Li, H., Liu, P., Xu, S., Li, Y., Dekker, J. D., Li, B., Fan, Y., Zhang, Z., Hong, Y., & Yang, G. (2017). FOXP1 controls mesenchymal stem cell commitment and senescence during skeletal aging. The Journal of Clinical Investigation, 127(4), 1241–1253.

López-Otín, C., Blasco, M. A., Partridge, L., Serrano, M., & Kroemer, G. (2023). Hallmarks of aging: An expanding universe. Cell.

Lu, A. T., Fei, Z., Haghani, A., Robeck, T. R., Zoller, J., Li, C., Lowe, R., Yan, Q., Zhang, J., & Vu, H. (2023). Universal DNA methylation age across mammalian tissues. Nature Aging, 3(9), 1144–1166.

Mace, G. M., Collar, N. J., Gaston, K. J., Hilton-Taylor, C., Akçakaya, H. R., Leader-Williams, N., Milner-Gulland, E. J., & Stuart, S. N. (2008). Quantification of extinction risk: IUCN’s system for classifying threatened species. Conservation Biology, 22(6), 1424–1442.

Maizels, R. M., & McSorley, H. J. (2016). Regulation of the host immune system by helminth parasites. Journal of Allergy and Clinical Immunology, 138(3), 666–675.

Marioni, R. E., Harris, S. E., Shah, S., McRae, A. F., von Zglinicki, T., Martin-Ruiz, C., Wray, N. R., Visscher, P. M., & Deary, I. J. (2016). The epigenetic clock and telomere length are independently associated with chronological age and mortality. International Journal of Epidemiology, 45(2), 424–432.

McLean, C. Y., Bristor, D., Hiller, M., Clarke, S. L., Schaar, B. T., Lowe, C. B., Wenger, A. M., & Bejerano, G. (2010). GREAT improves functional interpretation of cis-regulatory regions. Nature Biotechnology, 28(5), 495–501.

McNew, S. M., Boquete, M. T., Espinoza-Ulloa, S., Andres, J. A., Wagemaker, N. C., Knutie, S. A., Richards, C. L., & Clayton, D. H. (2021). Epigenetic effects of parasites and pesticides on captive and wild nestling birds. Ecology and Evolution, 11(12), 7713–7729.

Murray, M. H., Becker, D. J., Hall, R. J., & Hernandez, S. M. (2016). Wildlife health and supplemental feeding: A review and management recommendations. Biological Conservation, 204, 163–174.

Noyes, H., Stevens, J., Teixeira, M., Phelan, J., & Holz, P. (1999). A nested PCR for the ssrRNA gene detects Trypanosoma binneyi in the platypus and Trypanosoma sp. In wombats and kangaroos in Australia1. International Journal for Parasitology, 29(2), 331–339.

Pannella, G. (1971). Fish otoliths: Daily growth layers and periodical patterns. Science, 173(4002), 1124–1127.

Parsons, K. M., Haghani, A., Zoller, J. A., Lu, A. T., Fei, Z., Ferguson, S. H., Garde, E., Hanson, M. B., Emmons, C. K., & Matkin, C. O. (2023). DNA methylation-based biomarkers for ageing long-lived cetaceans. Molecular Ecology Resources, 23(6), 1241–1256.

Pastar, I., Marjanovic, J., Stone, R. C., Chen, V., Burgess, J. L., Mervis, J. S., & Tomic-Canic, M. (2021). Epigenetic regulation of cellular functions in wound healing. Experimental Dermatology, 30(8), 1073–1089.

Pearce, L. R., Huang, X., Boudeau, J., Pawłowski, R., Wullschleger, S., Deak, M., Ibrahim, A. F., Gourlay, R., Magnuson, M. A., & Alessi, D. R. (2007). Identification of Protor as a novel Rictor-binding component of mTOR complex-2. Biochemical Journal, 405(3), 513–522.

Pearson, J. C., Lemons, D., & McGinnis, W. (2005). Modulating Hox gene functions during animal body patterning. Nature Reviews Genetics, 6(12), 893–904.

Pedersen, A. B., & Fenton, A. (2015). The role of antiparasite treatment experiments in assessing the impact of parasites on wildlife. Trends in Parasitology, 31(5), 200–211.

Pepke, M. L. (2024). Telomere length is not a useful tool for chronological age estimation in animals. BioEssays, 46(2), 2300187.

Perna, L., Zhang, Y., Mons, U., Holleczek, B., Saum, K.-U., & Brenner, H. (2016). Epigenetic age acceleration predicts cancer, cardiovascular, and all-cause mortality in a German case cohort. Clinical Epigenetics, 8, 1–7.

Peters, A., Delhey, K., Nakagawa, S., Aulsebrook, A., & Verhulst, S. (2019). Immunosenescence in wild animals: Meta-analysis and outlook. Ecology Letters, 22(10), 1709–1722.

Peters, K. J., Gerber, L., Scheu, L., Cicciarella, R., Zoller, J. A., Fei, Z., Horvath, S., Allen, S. J., King, S. L., & Connor, R. C. (2023). An epigenetic DNA methylation clock for age estimates in Indo-Pacific bottlenose dolphins (Tursiops aduncus). Evolutionary Applications, 16(1), 126–133.

Pinho, G. M., Martin, J. G., Farrell, C., Haghani, A., Zoller, J. A., Zhang, J., Snir, S., Pellegrini, M., Wayne, R. K., & Blumstein, D. T. (2022). Hibernation slows epigenetic ageing in yellow-bellied marmots. Nature Ecology & Evolution, 6(4), 418–426.

Power, M. L., Ransome, R. D., Riquier, S., Romaine, L., Jones, G., & Teeling, E. C. (2023). Hibernation telomere dynamics in a shifting climate: Insights from wild greater horseshoe bats. Proceedings of the Royal Society B, 290(2008), 20231589.

Quach, A., Levine, M. E., Tanaka, T., Lu, A. T., Chen, B. H., Ferrucci, L., Ritz, B., Bandinelli, S., Neuhouser, M. L., & Beasley, J. M. (2017). Epigenetic clock analysis of diet, exercise, education, and lifestyle factors. Aging (Albany NY), 9(2), 419.

R Core Team. (2025). R: A Language and Environment for Statistical Computing [Computer software]. R Foundation for Statistical Computing. https://www.R-project.org/

Razin, A., & Kantor, B. (2005). DNA methylation in epigenetic control of gene expression. Epigenetics and Chromatin, 151–167.

Ritchie, M. E., Phipson, B., Wu, D., Hu, Y., Law, C. W., Shi, W., & Smyth, G. K. (2015). Limma powers differential expression analyses for RNA-sequencing and microarray studies. Nucleic Acids Research, 43(7), e47–e47.

Robeck, T. R., Haghani, A., Fei, Z., Lindemann, D. M., Russell, J., Herrick, K. E., Montano, G., Steinman, K. J., Katsumata, E., & Zoller, J. A. (2023). Multi-tissue DNA methylation aging clocks for sea lions, walruses and seals. Communications Biology, 6(1), 359.

Rowe, F., Bradfield, A., Quy, R., & Swinney, T. (1985). Relationship between eye lens weight and age in the wild house mouse (Mus musculus). Journal of Applied Ecology, 55–61.

Ryan, C. P., Lee, N. R., Carba, D. B., MacIsaac, J. L., Lin, D. T., Atashzay, P., Belsky, D. W., Kobor, M. S., & Kuzawa, C. W. (2024). Pregnancy is linked to faster epigenetic aging in young women. Proceedings of the National Academy of Sciences, 121(16), e2317290121.

Sagonas, K., Meyer, B. S., Kaufmann, J., Lenz, T. L., Häsler, R., & Eizaguirre, C. (2020). Experimental parasite infection causes genome-wide changes in DNA methylation. Molecular Biology and Evolution, 37(8), 2287–2299.

Schwanz, L. E. (2008). Chronic parasitic infection alters reproductive output in deer mice. Behavioral Ecology and Sociobiology, 62(8), 1351–1358.

Scott, M. (2023). Helminth-host-environment interactions: Looking down from the tip of the iceberg. Journal of Helminthology, 97, e59.

Sepers, B., Erven, J. A., Gawehns, F., Laine, V. N., & van Oers, K. (2021). Epigenetics and early life stress: Experimental brood size affects DNA methylation in great tits (Parus major). Frontiers in Ecology and Evolution, 9, 609061.

Shanebeck, K. M., Besson, A. A., Lagrue, C., & Green, S. J. (2022). The energetic costs of sub-lethal helminth parasites in mammals: A meta-analysis. Biological Reviews, 97(5), 1886–1907.

Shukla, V., Rao, M., Zhang, H., Beers, J., Wangsa, D., Wangsa, D., Buishand, F. O., Wang, Y., Yu, Z., & Stevenson, H. S. (2017). ASXL3 is a novel pluripotency factor in human respiratory epithelial cells and a potential therapeutic target in small cell lung cancer. Cancer Research, 77(22), 6267–6281.

Snir, S., Farrell, C., & Pellegrini, M. (2019). Human epigenetic ageing is logarithmic with time across the entire lifespan. Epigenetics, 14(9), 912–926.

Sullivan, I. R., Adams, D. M., Greville, L. J., Faure, P. A., & Wilkinson, G. S. (2022). Big brown bats experience slower epigenetic ageing during hibernation. Proceedings of the Royal Society B, 289(1980), 20220635.

Sweeny, A. R., Albery, G. F., Venkatesan, S., Fenton, A., & Pedersen, A. B. (2021). Spatiotemporal variation in drivers of parasitism in a wild wood mouse population. Functional Ecology, 35(6), 1277–1287.

Sweeny, A. R., Clerc, M., Pontifes, P. A., Venkatesan, S., Babayan, S. A., & Pedersen, A. B. (2021). Supplemented nutrition decreases helminth burden and increases drug efficacy in a natural host–helminth system. Proceedings of the Royal Society B, 288(1943), 20202722.

Telfer, S., Bown, K., Sekules, R., Begon, M., Hayden, T., & Birtles, R. (2005). Disruption of a host-parasite system following the introduction of an exotic host species. Parasitology, 130(6), 661–668.

Teschendorff, A. E., & Horvath, S. (2025). Epigenetic ageing clocks: Statistical methods and emerging computational challenges. Nature Reviews Genetics, 26(5), 350–368.

Turbill, C., Smith, S., Deimel, C., & Ruf, T. (2012). Daily torpor is associated with telomere length change over winter in Djungarian hamsters. Biology Letters, 8(2), 304–307.

van Iterson, M., van Zwet, E. W., Bios Consortium, & Heijmans, B. T. (2017). Controlling bias and inflation in epigenome-and transcriptome-wide association studies using the empirical null distribution. Genome Biology, 18(1), 19.

Walton, J. B., & Andrews, J. (1981). Torpor induced by food deprivation in the wood mouse Apodemus sylvaticus. Journal of Zoology, 194(2), 260–263.

Watson, H., Powell, D., Salmón, P., Jacobs, A., & Isaksson, C. (2021). Urbanization is associated with modifications in DNA methylation in a small passerine bird. Evolutionary Applications, 14(1), 85–98.

Wolf, S. E., Babayan, S. A., Marioni, R. E., Horvath, S., Pedersen, A. B., & Little, T. J. (2026). Data from: Epigenetic Clocks Uncover the Natural History of Ageing in Wild Mice [Dataset]. Zenodo. 10.5281/zenodo.20554783

Yu, G., Wang, L.-G., & He, Q.-Y. (2015). ChIPseeker: An R/Bioconductor package for ChIP peak annotation, comparison and visualization. Bioinformatics, 31(14), 2382–2383.

Zhao, L., Liu, D., Xu, J., Wang, Z., Chen, Y., Lei, C., Li, Y., Liu, G., & Jiang, Y. (2018). The framework for population epigenetic study. Briefings in Bioinformatics, 19(1), 89–100.

Zipple, M. N., Zhao, I., Kuo, D. C., Lee, S. M., Sheehan, M. J., & Zhou, W. (2025). Ecological Realism Accelerates Epigenetic Aging in Mice. Aging Cell, e70098.

Zoller, J. A., Parasyraki, E., Lu, A. T., Haghani, A., Niehrs, C., & Horvath, S. (2024). DNA methylation clocks for clawed frogs reveal evolutionary conservation of epigenetic aging. GeroScience, 46(1), 945–960.

