## Supplemental for "Epigenetic Clocks Uncover the Natural History of Ageing in Wild Mice"

**SUPPLEMENTARY MATERIAL for Epigenetic Clocks Uncover the Natural History of Ageing in Wild Mice**

| **Table S1.** Sample size of unique mice treated per treatment combination and for whom we have DNAm data. *Multi-Year mice were those who survived over the winter to be recaptured in 2024. | | | | | | | | | | | |
| --- | --- | --- | --- | --- | --- | --- | --- | --- | --- | --- | --- |
|  | 2023 | | | | | | 2024 | | | |  |
| Number Timepoints | 2 | | 1 | | 2+* | | 2 | | 1 | |  |
| **Treatment Combination** | **F** | **M** | **F** | **M** | **F** | **M** | **F** | **M** | **F** | **M** | **Total** |
| Control-Anthelminthic | 4 | 8 | 3 | 5 | 1 | 0 | 3 | 5 | 4 | 0 | 33 |
| Control-Control | 7 | 8 | 7 | 1 | 2 | 0 | 1 | 3 | 2 | 1 | 32 |
| Supplement-Anthelminthic | 15 | 14 | 7 | 2 | 1 | 1 | 1 | 3 | 1 | 4 | 49 |
| Supplement-Control | 9 | 12 | 6 | 9 | 1 | 1 | 1 | 2 | 2 | 6 | 49 |
| Total | 77 | | 40 | | 7 | | 19 | | 20 | | 163 |

**Figure S1**. Performance of the wood mouse epigenetic clock on fully independent test individuals (n = 14). (A) Predicted age from the wood mouse clock versus chronological age, showing median absolute error (MdAE) and correlation comparable to analyses using all test samples. (B) Predicted age from the pan-mammalian clock for the same individuals, showing lower accuracy than the wood mouse-specific clock.

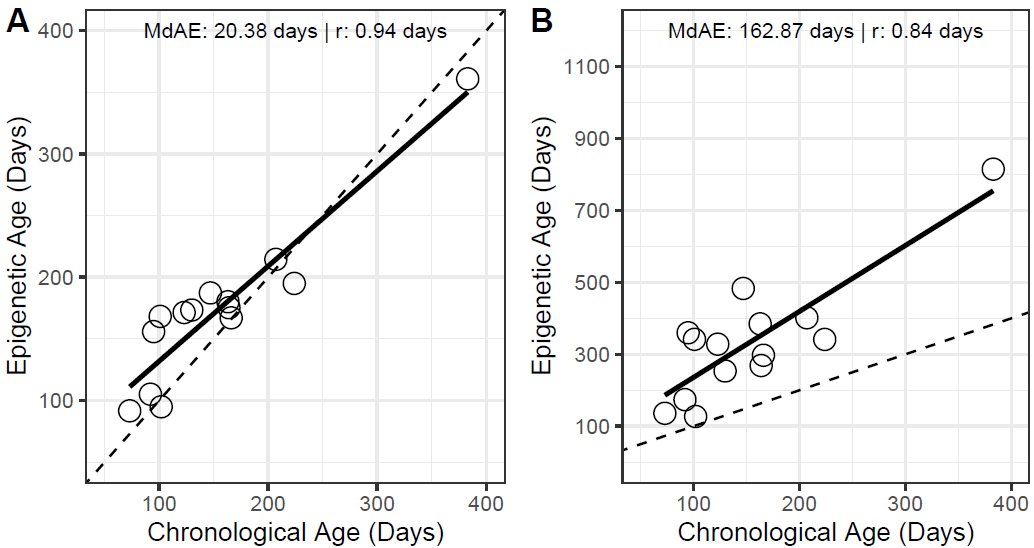

**Figure S2**. **Estimation of laboratory-reared wood mouse chronological age using three pan-mammalian clocks** (Lu et al., 2023). These clocks predict (A) log-transformed chronological age, (B) relative age (relative to the species maximum lifespan), and (C) log-linear age (relative to age at sexual maturity and gestation time). Each output was back transformed into age in days. Values for maximum lifespan, sexual maturity, and gestation were 6.3, 0.19, and 0.06 years, respectively. The dashed lines depict y = x.

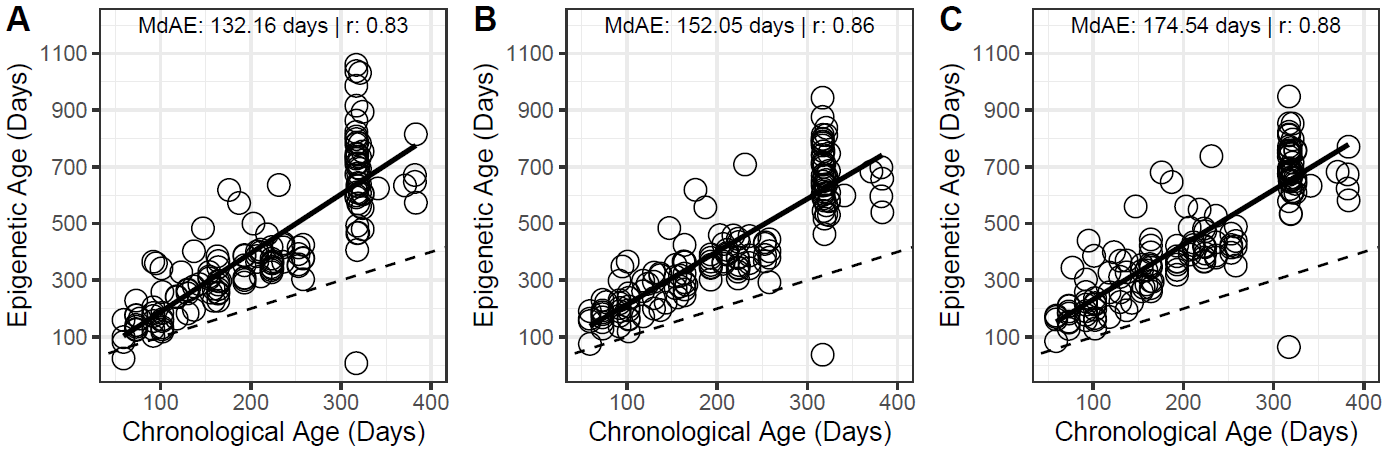

**Figure S3. Estimation of wild wood mouse chronological age using three pan-mammalian clocks** (Lu et al., 2023; see Figure S1 for descriptions of each). Because exact age is unknown, age estimates are regressed against age estimates generated from our wood mouse epigenetic clock. The dashed lines depict y = x.

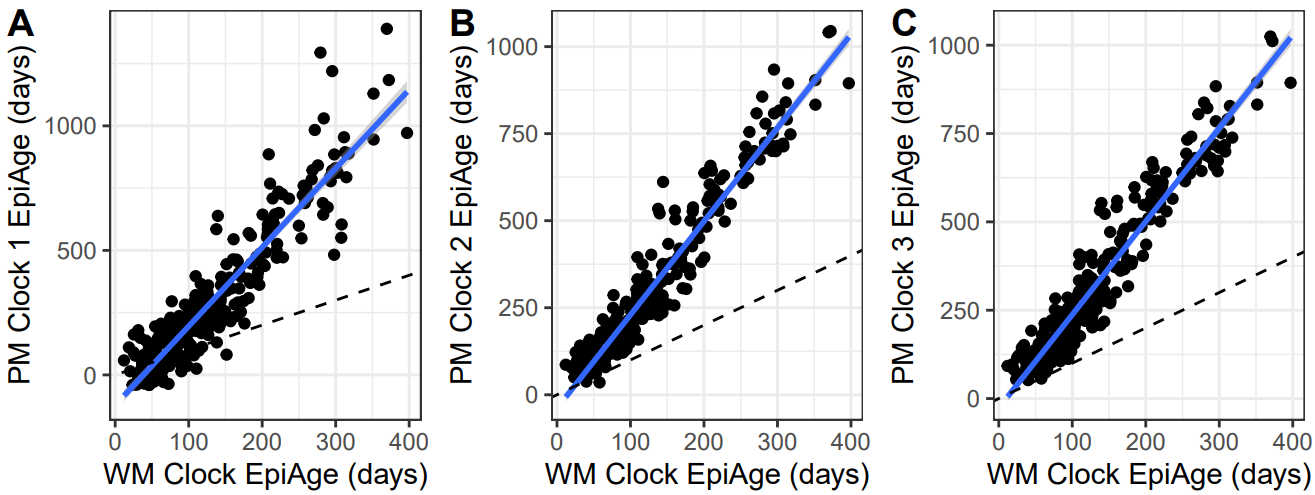

| **Table S2**. Epigenetic age significantly predicts the parasite diversity detected in a single day’s sample. Parasite diversity ranges from 0 to 7. | | | | |
| --- | --- | --- | --- | --- |
| **Predictor** | **Estimate** | **Std. Error** | **z value** | **Pr(>\|z\|)** |
| (Intercept) | 0.473453 | 0.427394 | 1.108 | 0.2680 |
| Food Supplement | -0.028203 | 0.117464 | -0.240 | 0.8103 |
| Epigenetic Age | 0.001951 | 0.000790 | 2.470 | 0.0135 * |
| Julian Date | 0.242391 | 0.521467 | 0.465 | 0.6421 |
| Sex (male) | 0.191473 | 0.131546 | 1.456 | 0.1455 |
| Reproductive Status | 0.232955 | 0.157524 | 1.479 | 0.1392 |
| Year (2024) | -0.232507 | 0.146227 | -1.590 | 0.1118 |

| **Table S3**. Summary of MANOVA predicting a multivariate outcome of 7 parasites with multiple fixed effects. | | | | |
| --- | --- | --- | --- | --- |
| **Predictor** | **Pillai** | **Approx F** | **DF** | **p-value** |
| Epigenetic Age | 0.3728 | 8.6607 | 7, 102 | <0.0001* |
| Supplement Tmt | 0.0239 | 0.3574 | 7, 102 | 0.9247 |
| Julian Date | 0.2079 | 3.8252 | 7, 102 | 0.0010* |
| Sex | 0.1475 | 2.5206 | 7, 102 | 0.0197* |
| Reproductive Status | 0.0672 | 1.0490 | 7, 102 | 0.4021 |
| Morphological Age | 0.0382 | 0.5794 | 7, 102 | 0.7713 |
| Year | 0.1017 | 1.6503 | 7, 102 | 0.1298 |

| **Table S4**. Post-hoc univariate tests following MANOVA analysis of predictors on multivariate parasite responses. All p-values were FDR adjusted for multiple comparisons. | | | |
| --- | --- | --- | --- |
| **Number of Ticks** | **F-value** | **p-value** | **p_adj** |
| Epigenetic Age | 18.8863 | 0.0000 | 0.0006* |
| Supplement Tmt | 0.6427 | 0.4245 | 0.7932 |
| Julian Date | 18.4782 | 0.0000 | 0.0006* |
| Sex | 2.8694 | 0.0932 | 0.4474 |
| Reproductive Status | 2.0237 | 0.1577 | 0.5153 |
| Morphological Age | 0.0059 | 0.9387 | 0.9787 |
| Year | 1.0234 | 0.3140 | 0.7326 |
| **Number of Mites** | **F-value** | **p-value** | **p_adj** |
| Epigenetic Age | 1.2286 | 0.2701 | 0.6975 |
| Supplement Tmt | 0.0848 | 0.7715 | 0.9105 |
| Julian Date | 0.0284 | 0.8664 | 0.9435 |
| Sex | 4.0338 | 0.0471 | 0.2885 |
| Reproductive Status | 0.0318 | 0.8589 | 0.9435 |
| Morphological Age | 0.1251 | 0.7243 | 0.8873 |
| Year | 3.5724 | 0.0614 | 0.3344 |
| **Number of Fleas** | **F-value** | **p-value** | **p_adj** |
| Epigenetic Age | 0.2829 | 0.5959 | 0.8456 |
| Supplement Tmt | 1.2963 | 0.2574 | 0.6975 |
| Julian Date | 8.5802 | 0.0041 | 0.0508* |
| Sex | 6.6302 | 0.0114 | 0.1115 |
| Reproductive Status | 0.0000 | 0.9975 | 0.9975 |
| Morphological Age | 0.5890 | 0.4445 | 0.7932 |
| Year | 0.7486 | 0.3888 | 0.7932 |
| **Bartonella Presence** | **F-value** | **p-value** | **p_adj** |
| Epigenetic Age | 5.7191 | 0.0185 | 0.1296 |
| Supplement Tmt | 0.4896 | 0.4856 | 0.7932 |
| Julian Date | 0.3921 | 0.5325 | 0.8154 |
| Sex | 2.5232 | 0.1151 | 0.4474 |
| Reproductive Status | 1.8722 | 0.1741 | 0.5331 |
| Morphological Age | 0.7424 | 0.3908 | 0.7932 |
| Year | 6.1413 | 0.0148 | 0.1205 |
| **WM Herpes Virus Presence** | **F-value** | **p-value** | **p_adj** |
| Epigenetic Age | 38.9236 | 0.0000 | <0.0001* |
| Supplement Tmt | 0.0087 | 0.9258 | 0.9787 |
| Julian Date | 1.2271 | 0.2704 | 0.6975 |
| Sex | 1.1235 | 0.2915 | 0.7143 |
| Reproductive Status | 0.5489 | 0.4604 | 0.7932 |
| Morphological Age | 0.2705 | 0.6040 | 0.8456 |
| Year | 0.4542 | 0.5018 | 0.7932 |
| **Trypanosoma Presence** | **F-value** | **p-value** | **p_adj** |
| Epigenetic Age | 0.1954 | 0.6593 | 0.8724 |
| Supplement Tmt | 0.1750 | 0.6766 | 0.8724 |
| Julian Date | 0.5809 | 0.4476 | 0.7932 |
| Sex | 0.0007 | 0.9793 | 0.9975 |
| Reproductive Status | 2.6030 | 0.1096 | 0.4474 |
| Morphological Age | 2.2917 | 0.1330 | 0.4654 |
| Year | 0.1355 | 0.7135 | 0.8873 |
| **H poly Egg Burden** | **F-value** | **p-value** | **p_adj** |
| Epigenetic Age | 0.4639 | 0.4972 | 0.7932 |
| Supplement Tmt | 0.0342 | 0.8536 | 0.9435 |
| Julian Date | 0.0781 | 0.7804 | 0.9105 |
| Sex | 2.4737 | 0.1187 | 0.4474 |
| Reproductive Status | 0.3156 | 0.5754 | 0.8456 |
| Morphological Age | 0.5812 | 0.4475 | 0.7932 |
| Year | 0.2306 | 0.6321 | 0.8603 |

**Experimental Treatments Methods and Analyses**

***Assessing general treatment effects of food supplementation and anthelminthic treatment in wild wood mouse populations***

*Analyses*: We assessed the effects of food supplementation and anthelminthic treatment on metrics related to reproduction and body condition. First, abundance of *Apodemus* was measured as the **minimum number known alive** (MNKA) for each of the four treatment grids during trapping sessions denoted as integers 1 – 9 (e.g., May to November, following Wolff 1996). We used a Poisson generalized linear model (GLM). Fixed effects included session, food supplementation treatment, and their interaction, as well as year and site. Second, we used a generalized linear mixed-effects model (GLMM) with a binomial error structure to examine the effects of experimental treatments and covariates on the **probability of being reproductively active**. The model included fixed effects for food supplementation, anthelminthic treatment, sex, their interaction (supplementation × sex), as well as scaled Julian date, site, and year. A random intercept for mouse ID accounted for repeated measures. The model was fit using the glmer function with the "bobyqa" optimizer and a high maximum iteration limit to ensure convergence. Last, we tested for effects on **mouse body condition** by fitting linear mixed-effects models with three response variables: **residual body mass index (rindex), raw body mass, and skeletal muscular index (SMI).** Each model included the fixed effects of anthelminthic treatment, food supplementation, reproductive status, sex, site, age class, scaled Julian date, and year, with mouse ID as a random effect. Significance of fixed effects was assessed using Type II Wald chi-square tests.

*Results*: Between 2023 and 2024, 491 unique wood mice were captured across our four grids and treated with either water or anthelminthic drug (see **Table S5** for sample sizes). The minimum number known alive (MNKA) increased over trapping sessions (*β* = 0.11 ± 0.02, *p* < 0.001), indicating rising grid-level abundance through the season. Food supplemented grids had significantly higher MNKA overall than controls (*β* = 0.40 ± 0.16, *p* = 0.014, **Figure S4**), but there was no evidence of a session-by-treatment interaction (*p* = 0.77). MNKA was lower in 2024 compared to 2023 (*β* = –0.92 ± 0.07, *p* < 0.001), and slightly higher at the Hewan site than at Penicuik (*β* = 0.20 ± 0.07, *p* = 0.002). While neither food supplementation nor anthelminthic treatment alone had significant effects on reproductive status (*p* = 0.30 and *p* = 0.17, respectively), mice in reproductive condition were significantly more likely to be found earlier in the season (*β* = –9.67 ± 0.99, *p* < 0.001), and were more likely to be male (*β* = 2.75 ± 0.42, *p* < 0.001), captured at the Penicuik site (*β* = 0.70 ± 0.25, *p* = 0.0057), and in 2024 (*β* = 1.46 ± 0.29, *p* < 0.001). The interaction between supplementation and sex was also not significant (*p* = 0.081). In addition, our treatments did not have an effect on residual body index or skeletal muscle index, although female, reproductive, adult individuals tended to have higher body condition-related indices (**Table S6**).

*Analyses*: We tested whether anthelmintic treatment reduced *H poly* infection intensity. *H poly* egg burden (eggs/gram faeces) was log(x+1)-transformed and rounded to the nearest integer. We used a zero-inflated negative binomial GLMM (glmmTMB), appropriate for the overdispersed and zero-inflated distribution of the response variable (mean = 2.57, variance = 5.48; ~40% zeros). Fixed effects included anthelminthic treatment (control, anthelminthic), food supplementation treatment (control, supplementation), scaled julian date, reproductive status (non-reproductive, reproductive), sex (male, female), site (Penicuik, Hewan), age (non-adult, adult), and year (2023, 2024), along with an interaction between anthelminthic treatment × first capture (to assess treatment impact before and after first treatments). Sampling date was formatted as Julian date, i.e., days since 1 Jan, and was scaled from 0 to 1 during modelling to reach model convergence. A random effect of mouse ID accounted for repeated measures. Zero inflation was modelled as a function of anthelminthic treatment. Post hoc pairwise comparisons for the anthelminthic × first capture interaction were performed using estimated marginal means (emmeans).

*Results*: We assessed 1176 faecal samples for *H poly* egg burden from 448 unique mice. Each mouse was sampled on average 2.6 ± 0.1 times from May to November (median = 2, range = 1 – 20 samples per mouse). Samples had an average eggs per gram faeces (EPG) of 126.0 ± 10.4 (median = 11.0 EPG, range = 0 – 5578 EPG). Prevalence of *H poly* ranged from 48% to 82% across groups. Before treatment, prevalence was relatively similar between Control and Anthelminthic groups (e.g., ~67% in 2023, ~50–53% in 2024). After treatment, prevalence was consistently higher in Controls (73–82%) compared to treated animals (65–73%), indicating a moderate reduction in infection prevalence following anthelmintic administration.

Anthelminthic treatment significantly reduced worm burdens, but only in individuals captured *after* treatment implementation (**Figure S5**). The interaction between anthelminthic treatment and first capture was significant (β = -0.193, SE = 0.092, *p* = 0.036), indicating that the treatment effect depended on capture timing. Post hoc contrasts showed no difference in worm intensity between treated and control animals caught before treatment (*p* = 0.79), but a significant reduction in treated animals after treatment compared to controls (β = -0.210, SE = 0.066, *p* = 0.0013). Food supplementation was also associated with a reduction in worm intensity (β = -0.124, SE = 0.046, *p* = 0.0066). Among covariates, infection intensity was slightly higher in 2024 compared to 2023 (β = 0.115, SE = 0.054, *p* = 0.033), while other predictors including sex, site, age, and reproductive status were not significant.

| **Table S5**. Sample size of mice per treatment combination. | | | | |
| --- | --- | --- | --- | --- |
| Year | Treatment | Total | Females | Males |
| 2023 | Control-Anthelminthic | 69 | 30 | 39 |
| 2023 | Control-Control | 70 | 35 | 35 |
| 2023 | Supplement-Anthelminthic | 95 | 47 | 49 |
| 2023 | Supplement-Control | 99 | 49 | 50 |
| 2024 | Control-Anthelminthic | 37 | 17 | 20 |
| 2024 | Control-Control | 35 | 14 | 21 |
| 2024 | Supplement-Anthelminthic | 44 | 20 | 24 |
| 2024 | Supplement-Control | 42 | 21 | 22 |

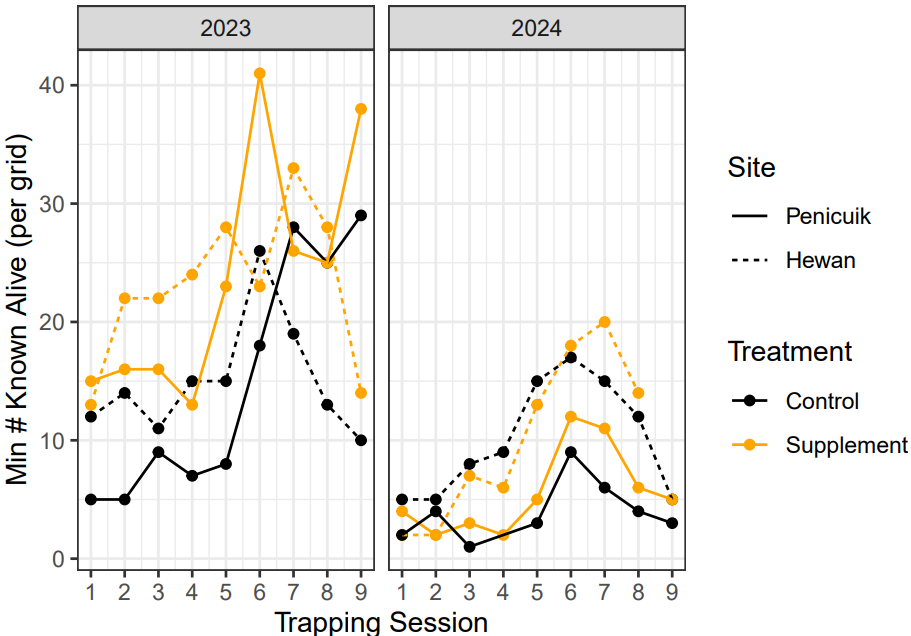

**Figure S4. Main treatment effect of food supplementation on wood mouse abundance**, measured as the minimum number known alive per each of four treatment grids from May to November in 2023-2024. Each grid was assigned the opposite food supplementation treatment between 2023 and 2024 replicates.

| **Table S6. Linear mixed-effects model of the effect of food supplementation and anthelminthic treatment on wood mouse body condition.** Mouse ID was included as a random effect. Test statistics are calculated using Type II sum of squares. Reference levels: no treatment with food or anthelminthic, and non-reproductive, male, and non-adult individuals. Sample sizes for analyses: body mass = 858 (from 455 mice); residual index and skeletal muscle index = 856 (from 454 mice). | | | |
| --- | --- | --- | --- |
| **Body Mass (g)** | | |  |
| **Predictor** | **β ± SE** | **t** | **p** |
| Food Supplement | 0.088 ± 0.243 | 0.36 | 0.718 |
| Anthelminthic | 0.243 ± 0.258 | 0.94 | 0.347 |
| Julian Date | -4.422 ± 0.869 | -5.09 | **<0.001^*^** |
| Reproduction Status | 3.618 ± 0.260 | 13.91 | **<0.001^*^** |
| Sex | -0.667 ± 0.272 | -2.45 | **0.015^*^** |
| Site | 0.031 ± 0.267 | 0.12 | 0.909 |
| Age Category | 4.529 ± 0.283 | 16.01 | **<0.001*** |
| Year | 0.963 ± 0.283 | 3.40 | **<0.001^*^** |
| **Residual Index** |  |  |  |
| **Predictor** | **β ± SE** | **t** | **p** |
| Food Supplement | -0.049 ± 0.216 | -0.23 | 0.820 |
| Anthelminthic | 0.126 ± 0.223 | 0.57 | 0.573 |
| Julian Date | -1.408 ± 0.805 | -1.75 | 0.081 |
| Reproduction Status | 2.084 ± 0.243 | 8.57 | **<0.001^*^** |
| Sex | -1.145 ± 0.235 | -4.86 | **<0.001^*^** |
| Site | -0.221 ± 0.229 | -0.96 | 0.338 |
| Age Category | 1.063 ± 0.261 | 4.07 | **<0.001^*^** |
| Year | -0.463 ± 0.248 | -1.87 | 0.063 |
| **Skeletal Muscle Index** | | |  |
| **Predictor** | **β ± SE** | **t** | **p** |
| Food Supplement | -0.070 ± 0.241 | -0.29 | 0.772 |
| Anthelminthic | 0.119 ± 0.246 | 0.49 | 0.628 |
| Julian Date | 0.657 ± 0.915 | 0.72 | 0.474 |
| Reproduction Status | 1.455 ± 0.277 | 5.26 | **<0.001^*^** |
| Sex | -1.271 ± 0.260 | -4.88 | **<0.001^*^** |
| Site | -0.276 ± 0.253 | -1.09 | 0.278 |
| Age Category | 0.215 ± 0.297 | 0.72 | 0.471 |
| Year | 1.433 ± 0.276 | 5.19 | **<0.001^*^** |
| ^1^Julian Date is scaled from 0 to 1 (end of May to November) | | | |

**Figure S5**. **The effect of food supplementation and anthelminthic treatment on *H poly* faecal egg burden in wood mice**, measured and plotted as raw log(x+1)-transformed eggs per gram faeces. Points represent individual samples.
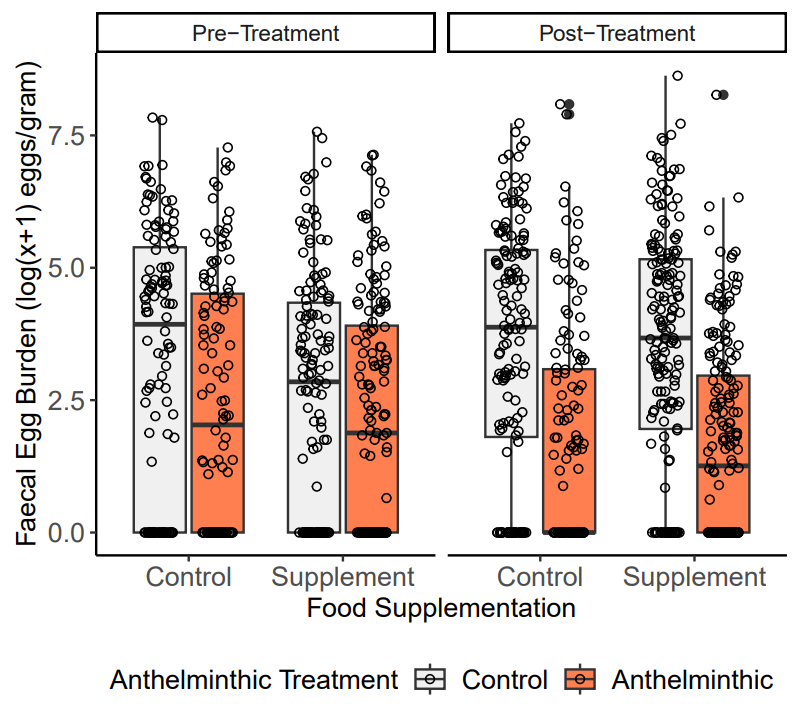

| **Table S7. Tukey-adjusted contrasts showing which of the four treatment combinations significantly differ in the change in epigenetic age over time.** | | | | |
| --- | --- | --- | --- | --- |
| **Contrast** | **β** | **SE** | **t** | **p** |
| (Control-Control)-(Control-Anthelminthic) | -0.04 | 0.26 | -0.15 | 0.99 |
| (Control-Control)-(Supplement-Control) | 0.09 | 0.23 | 0.37 | 0.98 |
| (Control-Control)-(Supplement-Anthelminthic) | 0.71 | 0.22 | 3.18 | 0.01 |
| (Control-Anthelminthic)-(Supplement-Control) | 0.12 | 0.24 | 0.51 | 0.96 |
| (Control-Anthelminthic)-(Supplement-Anthelminthic) | 0.75 | 0.24 | 3.19 | 0.01 |
| (Supplement-Control)-(Supplement-Anthelminthic) | 0.63 | 0.20 | 3.13 | 0.01 |

| **Table S8**. Predictors of the change in epigenetic age (difference) over a duration of time in wild wood mice. | | | |
| --- | --- | --- | --- |
| **Predictor** | **β ± SE** | **F value** | **P-value** |
| Combination Treatment | 52.94 ± 20.21 | 1.46 | 0.23 |
| Duration Between Samples | 0.90 ± 0.15 | 34.88 | < 0.0001 |
| Year | 2.05 ± 13.96 | 0.0007 | 0.98 |
| Sex (Male) | -4.54 ± 12.10 | 1.57 | 0.21 |
| Initial Reproductive Status (Yes) | -8.48 ± 12.33 | 1.33 | 0.25 |
| Combination x Duration | -0.41 ± 0.17 | 5.57 | 0.001 |

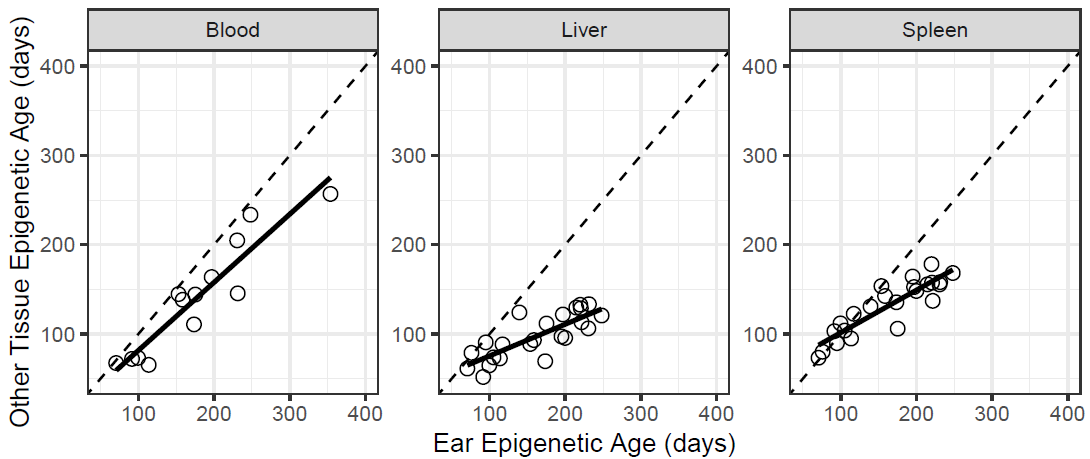
**Figure S5**. Correlation of predicted epigenetic age from ear biopsies compared to three other tissue types in known-age colony mice. All predicted epigenetic ages were calculated using the ear-specific wood mouse epigenetic clock developed in this manuscript. The dashed line indicates a perfect correlation between two tissues. When applied to these tissues, our epigenetic clock predicted actual age with these accuracies: Blood (MdAE = 24.4 days; corr = 0.935), Liver (MdAE = 65.2 days; corr = 0.83), and Spleen (MdAE = 30.2 days; corr = 0.89). As shown here, epigenetic age estimates were strongly correlated across tissues (Ear vs Blood: 0.93; Liver: 0.80; Spleen: 0.89), supporting the generality of the ear-based clock.
